# Mapping the gene space at single-cell resolution with gene signal pattern analysis

**DOI:** 10.1101/2023.11.26.568492

**Authors:** Aarthi Venkat, Sam Leone, Scott E. Youlten, Eric Fagerberg, John Attanasio, Nikhil S. Joshi, Michael Perlmutter, Smita Krishnaswamy

## Abstract

In single-cell sequencing analysis, several computational methods have been developed to map the cellular state space, but little has been done to map or create embeddings of the gene space. Here, we formulate the gene embedding problem, design tasks with simulated single-cell data to evaluate representations, and establish ten relevant baselines. We then present a graph signal processing approach we call *gene signal pattern analysis* (GSPA) that learns rich gene representations from single-cell data using a dictionary of diffusion wavelets on the cell-cell graph. GSPA enables characterization of genes based on their patterning on the cellular manifold. It also captures how localized or diffuse the expression of a gene is, for which we present a score called the *gene localization score*. We motivate and demonstrate the efficacy of GSPA as a framework for a range of biological tasks, such as capturing gene coexpression modules, condition-specific enrichment, and perturbation-specific gene-gene interactions. Then, we showcase the broad utility of gene representations derived from GSPA, including for cell-cell communication (GSPA-LR), spatial transcriptomics (GSPA-multimodal), and patient response (GSPA-Pt) analysis.

## 1 Introduction

A variety of techniques are used to map the cellular state space for single-cell RNA sequencing (scRNA-seq) analysis, including PCA, *t* -SNE, UMAP, HSNE, and PHATE [4, 6, 32, 69, 96]. These methods build low-dimensional embeddings based on the expression similarity between cells across thousands of genes. They reveal the organization of the cellular landscape, including clusters of similar cells, trajectories along phenotypic continuums, and multiscale structure fundamental to complex organisms. The expression of genes in cells is also highly organized. Genes are coordinated into complexes, biological processes, and pathways. However, despite the myriad techniques for understanding the cellular state space, it has not been possible to apply them analogously to understand the gene landscape. High and variable degrees of biological and technical noise [40], including “dropout”, or the lack of detection of an expressed gene due to sampling inefficiency [49], affect our ability to quantify gene-gene similarity and obscure the structure of the gene landscape.

Therefore, rather than building gene embeddings from single-cell data directly, we make the critical framing of genes as *signals* on a cell-cell graph, where the graph is commonly viewed as a discretization of an underlying cellular manifold. While graph feature or graph signal embeddings have been largely understudied compared to graph node embeddings in machine learning, the field of graph signal processing has made important advances by modeling features on graph nodes as signals or functions [72]. Analysis of such features extends classical signal processing concepts and tools, such as Fourier and wavelet analysis, to data residing on graphs.

We thus employ diffusion-based manifold learning for learning representations of signals on graphs. We first model random walks across the cell-cell graph and construct a diffusion operator **P**, which describes the transition probabilities of a single step between graph nodes. Powering **P** to *t* gives the transition probabilities of a *t*-step random walk, and thus *t* controls the scale of the representations; that is, small values of *t* capture local (smaller scale) representations and larger values capture global (larger scale) representations [21]. Based on these observations, we can leverage different values of *t* to produce multiscale representations of the cell-cell graph. Inspired by classical wavelet constructions [66], diffusion wavelets [22] power **P** to increasing scales and construct matrices defined by the differences between scales of diffusion (e.g., for two values to *t, t*_1_ and 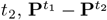). This allows them to play a powerful role in graph signal processing [10, 72, 81].

Here, we present gene signal pattern analysis (GSPA), a new method for embedding genes in single-cell datasets using a novel combination of diffusion wavelets and deep learning. First, we build a cell-cell graph (Figure 1a) to define genes measured as signals on the cell-cell graph (Figure 1b). Then, we decompose gene signals using a large dictionary of diffusion wavelets of varying scales centered on different vertices of the graph (Figure 1c). The result is a representation of each gene in a single-cell dataset as a set of graph diffusion wavelet coefficients. We pair this representation with an deep autoencoder framework to reduce the dimensionality of the gene representation and render it suitable for tasks such as visualization and prediction of gene characteristics (Figure 1d). In addition, we show the utility of ranking genes based on their distance to synthetic signals. This enables cell type association of gene signals, as well as a new type of analysis, which we term *differential localization* (Figure 1e). We highlight the importance of genes that are highly localized on the cellular manifold versus genes that are diffusely expressed, hypothesizing that localized genes are beneficial in characterizing different areas of the manifold informatively. As no previous tasks exist to evaluate algorithms for gene embeddings, we create benchmarking tasks based on three synthetic datasets with known ground truth. These tasks are designed to test the ability of gene embeddings to 1) preserve gene-gene relationships and 2) capture the gene localization score, which in turn enables 3) visualization of the gene space and 4) characterization of genes associated with gene modules, trajectories and archetypes. We also showcase these abilities on published datasets.

**Figure 1.**
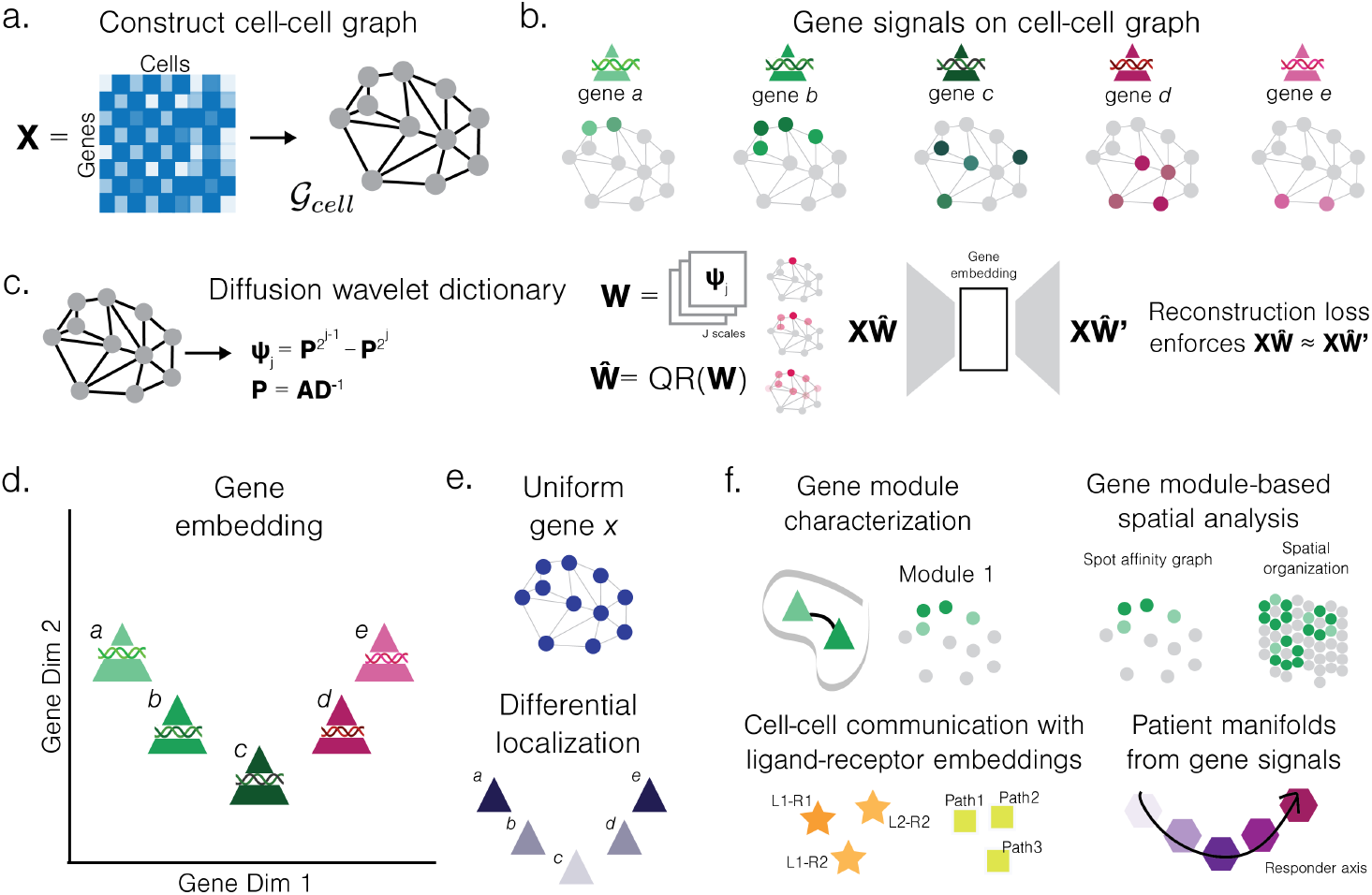
Overview of Gene Signal Pattern Analysis. a. Construction of cell-cell graph, where nodes are cells and edges are affinities between cells based on similarity of transcriptomic measurements. b. Five demonstrative gene signals (triangles), where signals are continuous functions defined on nodes of cell-cell graph. c. Construction of diffusion wavelets **W**, or QR-factorized dictionary 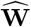 from scales 1, …, *J* to describe salient features of cell-cell graph. Gene signals are projected onto wavelet dictionary and gene embeddings learned via autoencoder architecture. d. Demonstrative gene embedding, where similar gene patterns are embedded closer together in the low-dimensional space, and far gene patterns are embedded far apart in the low-dimensional space. e. Differential localization determines how diffusely expressed gene signals are on graph, where very diffusely expressed signals do not explain cell-cell variation. *a* and *e* are most localized and *c* is least localized. f. Example downstream applications of gene signal pattern analysis, where gene embeddings enable cell type-independent characterization of gene modules, cell-cell communication, spatial transcriptomics, and patient manifolds. **P**: diffusion operator; **A**: adjacency/affinity matrix; **D**: degree matrix; L-R: ligand-receptor; Path: pathway

We demonstrate the utility of GSPA for biological interpretation in multiple settings (Figure 1f). First, we analyze a newly generated scRNA-seq dataset of CD8+ T cells in response to acute and chronic infection at days 4, 8, and 40. We visualize the gene space and show that localized genes correspond to meaningful biological processes. Additionally, we characterize gene modules and networks corresponding to different T cell programs, demonstrating GSPA is uniquely able to identify gene sets enriched for type 1 interferon signaling. We also identify gene networks perturbed in CD8+ T cells at acute infection day 8 in a newly generated single-cell CRISPR screen. This reveals gene-gene relationships involved in effector differentiation for *Tbx21* knockout and related to memory and effector capacity for *Klf2* knockout.

Importantly, the learned gene representations are highly informative and can be used for tasks beyond cellular heterogeneity characterization. To this end, we present a new approach for cell-cell communication analysis termed GSPA-based ligand-receptor analysis (GSPA-LR). By concatenating ligand and receptor embeddings into a single pair representation, then embedding ligand-receptor (LR) pairs, this approach identifies related LR patterns and pathways within and across multiple cell types. This is in contrast to most cell-cell communication tools, which are based on inferring communications between specific clusters or cell types. In an experimental model of peripheral tolerance [24], GSPA-LR analysis recovers condition-specific events in which multiple cell types activate signals for increased migration and effector activity in response to skin lesion, and a subset of myeloid cells sends signals to a subset of T cells via the PD-L1 checkpoint.

We also present a version of GSPA for multimodal data (GSPA-multimodal). This method uses a diffusion operator created jointly from different modalities, such as, for spatial transcriptomics data, gene expression and spatial coordinates. GSPA-multimodal identifies gene modules in key lymph node substructures corresponding to the blood vessel, germinal center, T cell zone, B follicle, B plasma cell region, and B interferon signature region [2]. We also highlight the relationship between differential localization and spatial variability and characterize microenvironmental signaling events between cell types enriched in each substructure.

Finally, we present a version of GSPA to map manifolds of patient samples (GSPA-Pt) which constructs patient vectors based on GSPA gene embeddings for improved prediction and interpretability. We demonstrate GSPA-Pt on 48 single-cell datasets from patients with melanoma pre- and post-immunotherapy [77], using these reduced representations to visualize the patient manifold and predict therapeutic response. GSPA-Pt identifies genes most predictive of response as associated with T cell progenitor states, and conversely genes predictive of non-response with T cell terminal differentiation states, consistent with prior literature.

Together, these results demonstrate the utility of considering gene expression measurements as signals on the cell-cell graph and leveraging fundamental ideas from graph signal processing for learning multiscale representations of signals on graphs. Code to run GSPA and generate results and figures here is available at https://github.com/KrishnaswamyLab/Gene-Signal-Pattern-Analysis.

## 2 Results

### 2.1 Gene embedding problem setup

We assume we are given a single-cell sequencing dataset consisting of *m* genes and their measurements in *n* cells, organized into an *m* × *n* matrix **X**. Here, the measured cells (columns of **X**) can be viewed as a sampling of possible cell states represented in the experiment and together make up the “cellular state space”. Despite the apparent high-dimensionality of this state space, cells are often modeled on an underlying manifold [43, 70, 99]. Manifold learning approaches have enabled myriad downstream tasks in single-cell analysis by modeling the constrained cellular state as having comparatively low intrinsic dimension. The majority of these approaches first build a cell-cell similarity graph 𝒢_*cell*_, where the vertices correspond to cells and edges correspond to cells with similar gene profiles. This graph represents a discretization of the cellular manifold (Methods).

Thus, with the insight that gene measurements may also be compressed into a lower-dimensional space for analyzing gene-gene relationships, our goal is to obtain a low-dimensional representation of each gene which preserves the inherent structure of the gene space with respect to the cellular manifold. In particular, we seek a reduced dimensional map *Θ* : ℝ^*n*^ →ℝ^*d*^, *d* ≪ *n*, which satisfies the desired properties enumerated below.

#### Desired Properties

1. **Preserving local and global distances between signals** A good gene embedding should produce similar representations of genes 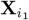 and 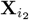 (viewed as rows of **X**) if they have similar measurement profiles. In order to ensure that we capture meaningful information, we aim to preserve distances based on the geometry of the underlying cell-cell graph 𝒢_*cell*_, rather than the naive pointwise distance, between gene signals.
2. **Noise robustness** Addressing biological noise, such as cell-to-cell variation, and technical noise, such as dropout, have been longstanding concerns in single-cell analysis and best practices [27, 40, 49, 65]. Due to variability in noise between genes with different expression levels [40], noise robustness is especially relevant for constructing gene embeddings. We thus seek a representation *Θ* such that 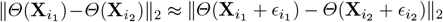 where 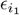 and 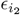 are measurement noise associated with genes 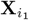 and 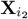.
3. **Flexibility to downstream tasks** Finally, we want to ensure our embedding *Θ* is flexibly defined for training on various additional tasks, whether concurrently with the learned embedding or downstream of the embedding. In the following section, we present our approach, gene signal pattern analysis (GSPA), to achieve these desired properties.

### 2.2 Gene signal pattern analysis (GSPA) overview

In order to construct the map *Θ*, we make the critical observation that the expression pattern **X**_*i*_ for gene *i* can be described as a signal (function) defined on the nodes of a cell-cell similarity graph 𝒢_*cell*_. Through this framing, we can compare how gene expression patterns are similar to and different from each other based on distances along the cellular manifold.

We first model random walks on the cell-cell similarity graph with a diffusion operator **P** that contains transition probabilities between cells on the basis of their similarity. We use this operator to construct a dictionary of diffusion wavelets for decomposing gene signals on the cell similarity matrix. In order to construct these wavelets, we power **P** to different values of *t* to capture local (small *t*) and global (large *t*) representations. Each diffusion wavelet is then defined by the difference of these powered diffusion operators ([22], Methods).

We then represent gene signals with respect to these local and global geometric structures. To this end, we decompose each signal using the wavelet dictionary, encoding the topology of the nodes (cells) it is defined on in addition to the signal (gene) itself. Finally, we learn a meaningful low-dimensional embedding using an autoencoder for reducing the wavelet-based representations (Algorithm 1). This reduced embedding can be used for many downstream analyses including visualization, gene module identification, or differential localization. We add further discussion and expand upon each step of this algorithm in the Methods section.

#### Algorithm 1

Gene signal pattern analysis algorithm for learning gene representations

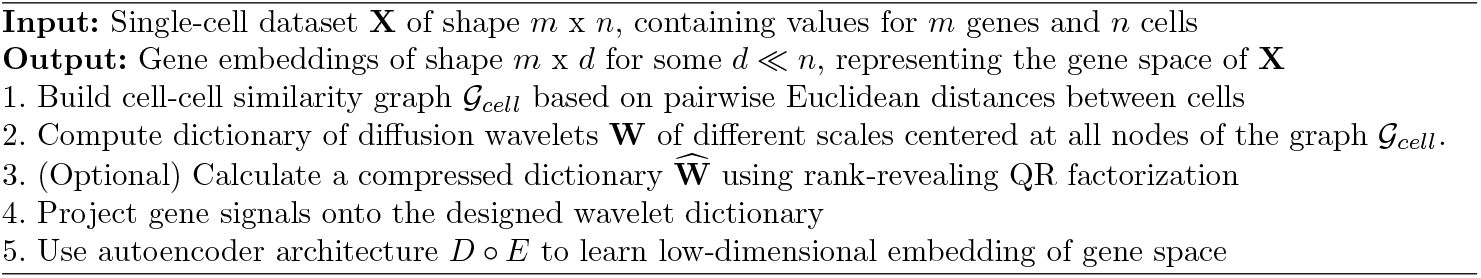

#### Constructing a cell-cell similarity graph from single-cell dat

In scRNA-seq, each cell is measured as a vector of gene expression counts. That is, the output of a scRNA-seq experiment is a matrix **X** of shape *m* x *n*, where *m* corresponds to the number of genes, and *n* the number of cells. The first step of the GSPA algorithm is to build a graph 𝒢_*cell*_ = (*V*_*cell*_, *E*_*cell*_), where each node in *V*_*cell*_ corresponds to a cell, and each edge 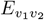 in *E*_*cell*_ describes the similarity between cell *v*_1_ and cell *v*_2_.

Due to the flexibility of graph construction, GSPA can handle a variety of use cases. Graphs can be constructed with affinities derived from multiple modalities, then used with GSPA for integrated gene analysis (Methods). Moreover, graphs can consist of cells from multiple sequencing runs. In such cases, downstream analysis is often affected by batch effects, in which expression of genes systematically differs between batches, resulting in cellular affinities defined by batch rather than cell type or other underlying biology. Batch effect affects all downstream analysis, including gene embeddings derived from our approach, where genes separate by enrichment within a particular batch (Extended Data Figure 1a). GSPA handles batch effect through either accepting batch-corrected cell measurements or cell embeddings as input, or correcting for batch in the cell-cell graph through mutual nearest neighbors (MNN) graph construction [42]. This results in gene embeddings corrected for differences in batch (Methods, Extended Data Figure 1b).

For large graphs, GSPA utilizes diffusion condensation, a coarse-graining process which iteratively condensing datapoints toward local centers of gravity [11, 57]. This technique allows GSPA to summarize the underlying topology of the data manifold in a smaller coarse-grained cell-cell graph for improved scalability (Methods, Extended Data Figure 10a).

#### Building dictionary of graph diffusion wavelets for gene representatio

Given the cellular graph 𝒢_*cell*_, we use data diffusion to model random walks across the cell-cell graph and uncover the underlying manifold. To facilitate these random walks, we define the diffusion operator **P** = **AD**^−1^, a row-normalized version of the affinity matrix **A** (with **D** denoting the degree matrix), where the entries of the diffusion operator correspond to the probability of moving from one cell to another (Methods). Powering the diffusion operator to the power of *t* corresponds to a *t*-step random walk through the data, and therefore each column of **P**^*t*^ may be viewed as a vector of weighted neighbors associated with each cell in our data.

To represent the multiscale structure of 𝒢_*cell*_, we power **P** to increasing scales and construct matrices defined by the difference of these powered diffusion operators, termed diffusion wavelets [22]. We construct diffusion wavelet of scale *j* centered at vertex *v* by 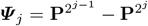 for 1 ≤ *j* ≤ *J* (and ***Ψ***_0_ = **I** − **P**) and extract the *v*-th row via 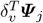, where *δ*_*v*_ is the Kronecker delta centered at the *v*-th vertex (Methods). These diffusion wavelets are then organized into a dictionary **W** of shape *n* x *Jn*. The number of scales for the wavelet dictionary *J* is defined as the *log* of the number of cells *n* based on Lemma 1 introduced and proven in [93] (Methods).

In general, the numerical rank of ***Ψ*** _*j*_ decreases as *j* increases, and furthermore, for large values of 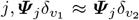 if vertex *v*_1_ is near vertex *v*_2_ on the graph. Therefore, a small set of large wavelets can be used to describe the graph at a coarse resolution. To reduce this redundancy, we leverage rank-revealing QR factorization as in [22], resulting in a compressed wavelet dictionary 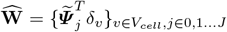 (Methods). We test gene signal embeddings both with (GSPA+QR) and without (GSPA) compression.

#### Projecting signals onto the wavelet dictionary

Each gene signal **X**_*i*_ of shape 1× *n* corresponds to the expression of the gene in the cellular state space. Importantly, we consider each measured gene feature as a signal on the cellular graph. Given a gene signal **X**_*i*_ and wavelet dictionary 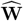 (or **W**), we project **X**_*i*_ onto the dictionary. For all gene signals, this corresponds to 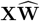 or **XW**. Theorem 1 stated below shows that the wavelet projection **X** → **XW** is continuous with respect to an Unbalanced Diffusion Earth Mover’s Distance (UDEMD) [91]. Intuitively, this UDEMD describes how similar two gene signals 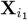 and 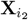 are in a manner informed by the geometry of the cellular graph (Methods).

#### Learning a low-dimensional representation with autoencoder for downstream analysis

As 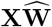 and **XW** correspond to a meaningful representation of genes, we can use this matrix for downstream analyses. However, this representation is unnecessarily high-dimensional, such that each feature corresponds to the gene signal projection of a particular cell at a particular diffusion scale. To reduce the dimensionality of the representation, we leverage an autoencoder *D* ○ *E*, consisting of an encoder *E* and decoder *D* such that *D* ○ *E* is approximately the identity (i.e.,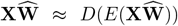), in order to learn a low-dimensional, information-rich gene embedding *E* 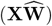 (Methods).

#### The GSPA architecture

Combining the above steps, we define gene signal pattern analysis (GSPA) by

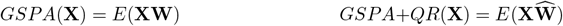

where **W** and 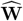 are uncompressed and compressed wavelet dictionaries (respectively), *E* is the encoder discussed above, and *GSPA* and *GSPA* + *QR* are taken to be the map *Θ* discussed in the problem setup (Methods).

### 2.3 Achieving desired representation properties with GSPA

#### Distance preservation

As discussed in Section 2.1, our first desired property is an embedding that preserves distances (quantified in a manner informed by the geometry of the cellular state space). Theorem 1 shows that GSPA is able to achieve this goal since it guarantees we will have *GSPA*(**X**_*i*_) ≈ *GSPA*(**X**_*j*_) whenever **X**_*i*_ is close to **X**_*j*_ with respect to the Unbalanced Diffusion Earth Mover’s Distance (UDEMD). This distance, a variant of traditional earth mover’s distance (EMD), views the signals **X**_*i*_ (when properly normalized) as probability distributions on the graph. Roughly speaking, UDEMD aims to compute the cost of rearranging one “mass distribution” into another, where the cost of moving mass from one cell to another is given by the length of the shortest graph path between the cells.

##### Theorem 1.

*The diffusion wavelet projection* **W** *is Lipschitz Continuous with respect to UDEMD, i*.*e*., *there exists a constant C >* 0 *such that*

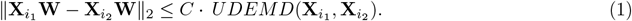

For a proof of Theorem 1 and a detailed statement and discussion of UDEMD, please see Section 6.3.

#### Noise robustness

Robustness to biological and technical noise is a key feature of diffusion-based single-cell analysis approaches [70]. Note that raising the diffusion operator **P** to the power *t* is equivalent to powering the eigenvalues of the diffusion operator by *t*, i.e., **P** = ***ΣΛΣ***^−1^, where the columns of ***Σ*** contain the (right) eigenvectors of **P** and ***Λ*** is a diagonal matrix whose entries are the corresponding eigenvalues (Methods). Thus, **P**^*t*^ = ***ΣΛ***^*t*^***Σ***^−1^ and powering **P** effectively results in powering the eigenvalues contained in ***Λ***. The eigenvectors are decreasingly ordered by their “frequency”, a notion of how rapidly a signal oscillates over the graph (Methods). It is known that 1 = *λ*_1_ *> λ*_2_≥ *λ*_3_ ≥….Therefore, powering **P** preserves the lead eigenspace and suppresses the subsequent spaces by a factor of 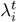.Acting on a signal **X**_*i*_ by **P**^*t*^ preserves the portion of the signal aligned with the first eigenvector and depresses the portion of the signal corresponding to the other eigenvectors by a factor of 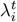.As *t* increases, the high-frequency (small eigenvalue) portion of the signal is suppressed. Naturally occurring signals tend to vary slowly and smoothly over the graph (and thus lie in the low-frequency eigenspaces), whereas noise is not related to the structure of the graph and therefore will often lie in the higher frequencies. In this manner, acting on the signal **X**_*i*_ by **P**^*t*^ has a denoising effect since it suppresses the high-frequency (noisy) portion of the signal. Therefore, we can restrict the dictionary to wavelets that decompose only the lower frequencies by initially multiplying each wavelet by **P**^*t*^. Additionally, we note that the distance preservation result Theorem 1, shows that the wavelet projection is continuous with respect to the unbalanced diffusion earth movers distance, which may be viewed as a form of noise robustness.

#### Flexibility to downstream tasks

Finally, we achieve flexibility by leveraging an autoencoder framework to learn a generalizable low-dimensional representation. In the following sections, we highlight this flexibility. First, we describe two gene rankings enabled by GSPA representations (Section 2.4). Then, we benchmark GSPA against various baselines for the preservation of gene-gene distances and gene localization on three simulated datasets (Section 2.5). We showcase GSPA for characterizing CD8+ T cell states (Section 2.6), then present three adaptations of gene signal pattern analysis for downstream tasks beyond characterizing cellular heterogeneity. GSPA-LR (Section 2.7) performs cluster-independent cell-cell communication analysis by constructing embeddings of ligands and receptors, then concatenating and embedding ligand-receptor pairs. This approach identifies shared patterns of ligand-receptor pairs on the cellular manifold without cell type annotation. GSPA-multimodal (Section 2.8) demonstrates the ability of GSPA to naturally integrate multiple modalities into a single gene embedding representation using integrated diffusion [56]. With spatial transcriptomics data, GSPA-multimodal characterizes gene programs with respect to both expression and spatial similarity. Finally, GSPA-Pt (Section 2.9) builds patient representations based on patient-specific gene embeddings, capturing both the topology of the cellular state space and gene-gene relationships for downstream prediction and exploration.

### 2.4 Capturing rankings based on embedded distances from synthetic signals

Beyond preserving relationships within the gene space, GSPA also preserves distances to other signals one can define on the shared cell-cell graph. This enables flexible ranking of genes based on distance to the synthetic signals. Here, we describe two approaches based on this idea for cell type association of gene signals and gene localization.

#### Cell type association of gene signals

In systems where cells naturally can be distinguished by cell types, it is standard to characterize cell types by differentially expressed genes [65]. Gene signal pattern analysis naturally enables identification of cell type-specific genes. Given a dataset where each cell is assigned a cluster (or annotated cluster, i.e. cell type), for each cell type *C*, we can define a set indicator signal **1**_*C*_ on all vertices of the cell-cell graph, where **1**_*C*_(*v*) = 1 if *v* ∈ *C* and **1**_*C*_(*v*) = 0, otherwise for *v* ∈ *V*_*cell*_. Then, we can rank genes in **X** based on how close they are to **1**_*C*_ in their dictionary representation. Formally, we define cell type association ranking of genes by the following score:

##### Definition 1.

*Given a normalized cell type indicator signal for cell type* 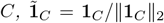*and wavelet representation* 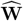, *the cell type association score, c*(*i*) *for each gene signal* **X**_*i*_ *is defined as:*

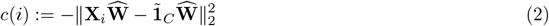

Unlike many differential expression tests, which compare mean expression between clusters, the cell type association score ranks highly genes that are close to zero expression in all other cell types and close to uniform expression in the cell type of interest. This results in a ranking that prioritizes specificity to the cell type and achieves close results to ground truth differential expression scores from simulated data (Extended Data Figure 1c). Additionally, conventional methods for detecting differential gene expression between clusters often return inflated p-values because of the double use of gene expression data, first to partition the data into clusters and then to define significance statistics along the same partitions [84]. Consequently, filtering based on p-value alone results in an increased rate of false positives, and some pipelines have turned to gene rankings instead of cutoffs [98].

#### Differential localization of gene signals on the cell-cell graph

For many single-cell datasets, including those that capture cellular trajectories, fine-grained cellular subtypes within coarse-grained cell types, rare cell types, and highly plastic cell types, it can be nontrivial to assign cluster labels and define cell state-specific genes [51]. Thus, in order to characterize cell-cell variation and differences between cell states, there is a need to identify genes specific to particular populations without prior clustering or annotation.

Addressing this need, we present an alternative strategy to calculate the specificity of a gene signal **X**_*i*_ across the cell-cell graph, termed differential localization. We make the observation that genes that are uniformly expressed across all cells are least likely to be involved in cell state-defining biological processes, in line with previous work [98]. Leveraging this insight, we calculate the distance between each gene signal and a multiscale representation of a uniform (constant) signal 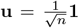, normalized in the same way we normalize other signals. We then rank genes by gene localization score *ℓ*, where genes most differentially localized are those farthest from the uniform signal representation. Formally,

##### Definition 2.

*The gene localization score, ℓ*(*i*) *for each gene signal* **X**_*i*_, *with normalized uniform signal* **u** *and wavelet representation* 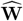, *is defined as:*

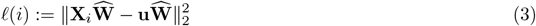

Genes with a high localization score are considered more relevant for describing cell-cell variation and can be used for feature selection or characterization of gene programs and networks without the underlying assumption of discrete clusters.

### 2.5 Comparison to alternative gene mapping strategies

We evaluate embeddings on three single-cell benchmarking datasets simulated with Splatter (one trajectory, two branches, and three branches) [106]. Using Splatter allows us to generate corresponding noisy gene expression data **X** and unseen true (noiseless) counts **X**^*′*^ (Figure 2a). By increasing the dropout probability to 95%, we ensure that the noisy gene expression data reflects levels of noise seen in real single-cell datasets. The linear trajectory dataset was 84.3% sparse, the two branch dataset was 85.3% sparse, and the three branch dataset was 85.3% sparse. We adapt ten baselines for comparisons, and we assess the embeddings for preservation of coexpression, fidelity of visualization, coherence of downstream analyses, and ability to capture signal localization.

**Figure 2.**
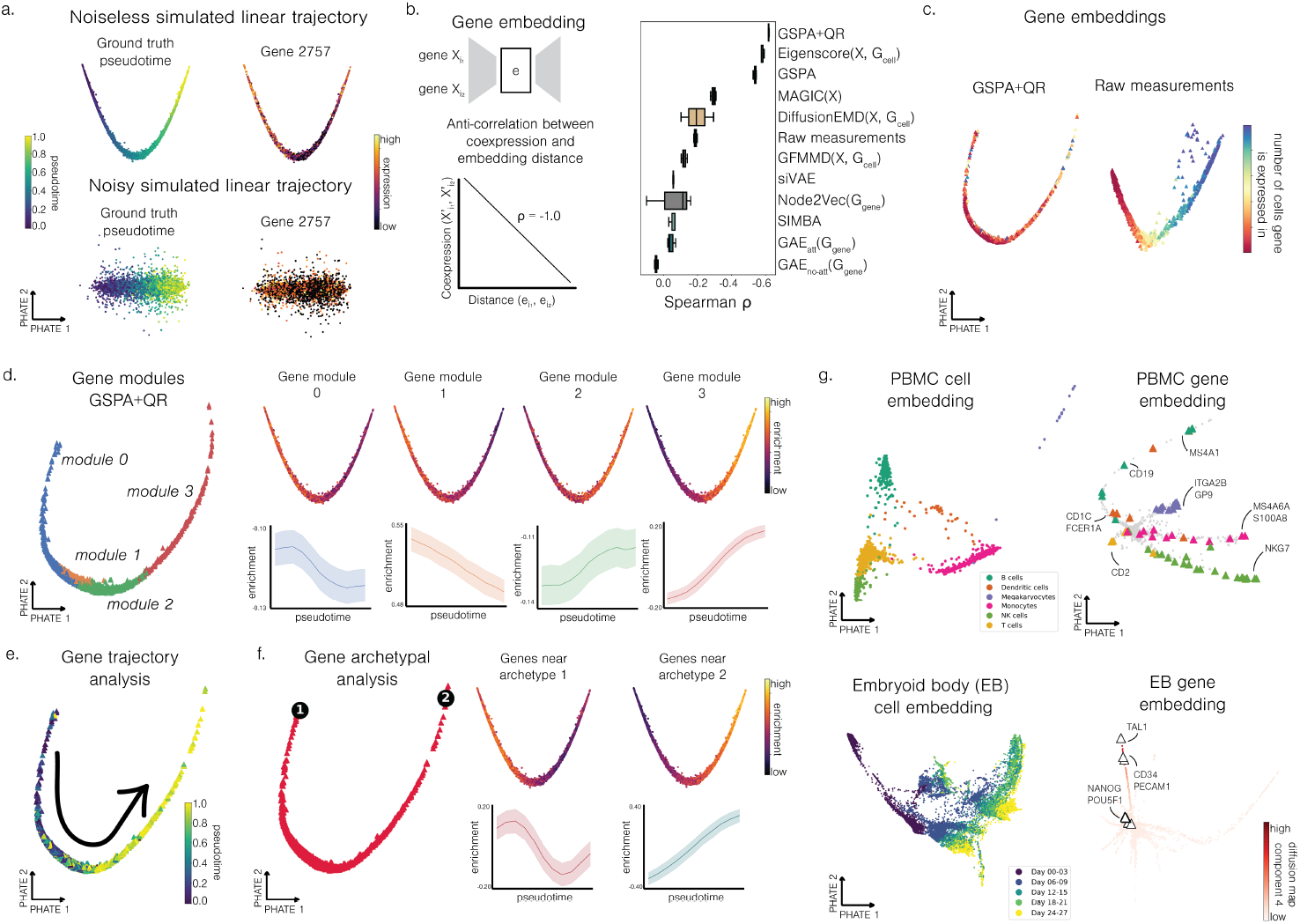
Gene signal pattern analysis preserves coexpression between genes and captures coherent visualization and gene modules, trajectories, and archetypes. a. Noiseless (top) and noisy (bottom) cell embeddings of simulated linear trajectory, colored by ground truth pseudotime provided by the simulation engine (left) and example gene expression (right). b. Experimental setup (left) and Spearman correlation evaluating performance on task for all comparisons across 3 runs (right). c. Gene embeddings of GSPA+QR and raw measurements colored by number of cells gene is expressed in. d. Gene modules detected by Leiden clustering (left) for GSPA+QR, and gene module enrichment and expression over time (right). e. GSPA+QR gene embedding colored by time at which gene peaks. f. GSPA+QR gene embedding with archetypes identified via AAnet (left), with gene enrichment and expression over time visualized for “archetypal” genes (genes closest to each archetype). g. PBMC cell embedding and gene embedding with key PBMC markers annotated from PanglaoDB (top) and EB cell embedding and gene embedding, colored by diffusion eigenvector and key hemangioblast lineage markers annotated from [69].

#### Description of gene mapping strategies adapted for comparison

For comparison, we pull from prior literature and design additional strategies based on methodology not previously used for gene embeddings. We summarize and diagram these comparisons in Extended Data Figure 2 and expand upon each approach in the Methods section.

First, we can map the gene space with the original measurements **X**, using an autoencoder to reduce the dimensionality. We term these embeddings the “raw” measurements. Alternatively, we can construct a gene-gene *k*-NN graph 𝒢_*gene*_ = (*V*_*gene*_, *E*_*gene*_) from **X** and employ three graph representation learning methods GAE_att_(𝒢_*gene*_) (a graph autoencoder with graph attention layers) [50], GAE_no-att_(𝒢_*gene*_) (a graph autoencoder with graph convolutional layers) and Node2Vec(𝒢_*gene*_) [39]. These approaches do not use the cell-cell graph to compute gene representations, which, as described in our first desired property (Section 2.1), may not sufficiently preserve distances. Therefore, we additionally compare against MAGIC [27], which denoises **X** via data diffusion on 𝒢_*cell*_, then use an autoencoder to reduce the dimensionality of the resulting denoised measurements. We demonstrate how diffusion wavelets differ from and improve upon the diffusion-based approach in MAGIC in Extended Data Figure 3, Extended Data Figure 4, and in the Methods section. We also compare against techniques that consider genes as signals on 𝒢_*cell*_. Diffusion Earth Mover’s Distance (DiffusionEMD) [93] computes optimal transport, or Wasserstein’s distance, based on multiscale diffusion kernels. Graph Fourier Mean Maximum Discrepancy (GFMMD) [62] is defined via an optimal witness function that is smooth on the graph and maximizes the difference in expectation between the pair of gene distributions. Furthermore, eigenscores [44], proposed for feature selection, were shown to be useful for mapping the gene space based on their alignment to low-frequency patterns in the data. We again reduce the dimensionality of these resulting multiscale gene representations via an autoencoder.

Finally, we consider two neural network approaches that are designed to jointly infer cell-wise and feature-wise embeddings. siVAE [19] leverages a cell-wise VAE and a feature-wise VAE, pre-training each before jointly training all model parameters for interpretable downstream analysis. SIMBA [15] co-embeds cells and features, through, for scRNA-seq analysis, constructing a heterogeneous graph of cells and genes (where an edge is added between a cell and gene if the gene is expressed in that cell) and learning an embedding of the graph. For both approaches, we train the entire model and analyze the resulting gene embeddings.

In summary, the baselines are as follows:

- Raw measurements approach embeds **X**.
- GAE_no-att_(𝒢_*gene*_) embeds 𝒢_*gene*_.
- GAE_att_(𝒢_*gene*_) embeds 𝒢_*gene*_.
- Node2Vec(𝒢_*gene*_) embeds 𝒢_*gene*_.
- MAGIC(**X**) embeds **X** after denoising with 𝒢_*cell*_.
- DiffusionEMD(**X**, 𝒢_*cell*_) embeds **X** via optimal transport on 𝒢_*cell*_.
- GFMMD(**X**, 𝒢_*cell*_) embeds **X** via MMD on 𝒢_*cell*_.
- Eigenscore(**X**, 𝒢 _*cell*_) embeds **X** via alignment to Laplacian eigenvectors of 𝒢_*cell*_.
- SIMBA co-embeds **X** and **X**^*T*^ via heterogeneous graph embedding.
- siVAE co-embeds **X** and **X**^*T*^ via jointly trained cell-wise and feature-wise VAEs.

These approaches represent a wide array of techniques adapted from graph signal processing, graph representation learning, and single-cell analysis for fair comparison.

#### Preservation of coexpression relationships in gene embeddings

For our first task for comparison, we recognized that genes that are coexpressed should be close in a low-dimensional embedding and genes that are expressed in very different cells should be far apart. To assess this, we first define the coexpression between genes *i*_1_ and *i*_2_ as the correlation of the true counts 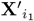 and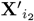. Then, we learn gene embeddings 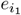 and 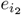 from the noisy counts **X** and compute their distance. Finally, we use the correlation between this distance and their coexpression as a metric for evaluation (Methods, Figure 2b, Extended Data Figure 5a). GSPA+QR performed best on all three benchmarking datasets, followed by GSPA on average (Figure 2b, Extended Data Figure 5b, Extended Data Figure 5c).

#### Visualization not impeded by technical factors of gene variation

We next assessed the ability to visualize the gene embeddings, which remains a useful tool for characterization and hypothesis generation in single-cell analysis [6, 69, 97]. We visualized each gene embedding with PHATE [69] and determined that, for the majority of methods, the major axis of variation captured by visualization is strongly correlated with the number of cells the gene is expressed in (Figure 2c, Extended Data Figure 6). This is despite normalizing each gene signal (Methods). By contrast, GSPA+QR and GSPA gene visualizations are not impeded by this confounding factor of gene expression.

#### Coherence of gene modules, trajectories, and archetypes on the cellular manifold

We next evaluated the coherence of downstream analyses derived from the gene embeddings. First, in all three datasets, we clustered the GSPA+QR gene embedding space to reveal groups of genes, termed gene modules. We then visualized the enrichment of each gene module on the cell embedding, and found that genes within each module show enriched activation in distinct regions of the embedding. We visualize these enrichment scores over cellular pseudotime and see that the gene modules share a related expression trend over time (Methods, Figure 2d, Extended Data Figure 5d, Extended Data Figure 5e). For comparison, we learn gene embeddings for all other approaches and identify gene modules. Across all other approaches, gene module enrichment scores are not interpretable and do not show a relationship with time (Extended Data Figure 7). We additionally show improvement over factor analysis method cNMF [54], which identifies gene expression programs (i.e. modules) but does not map the gene space for visualization or other analyses.

To highlight the importance of these other downstream analyses, we further explore the gene embeddings using trajectory analysis and archetypal analysis, two other approaches used for characterizing cellular relationships. Gene trajectory analysis shows that genes are organized by the pseudotime at which they peak in expression on the cellular trajectory (Figure 2e), suggesting that GSPA embeddings can reveal ordering of genes related to particular biological processes. Gene archetypal analysis with two archetypes reveals that genes closest to each archetype show enriched expression on each side of the trajectory and reflect an association with pseudotime (Figure 2f). Again, almost all comparisons do not capture these trends, suggesting they are not sufficiently capturing gene-gene relationships for downstream analysis (Extended Data Figure 7).

In publicly-available single-cell datasets, GSPA+QR identifies known relationships between genes (Methods, Figure 2g). For 3k peripheral blood mononuclear cells (PBMCs) from a healthy donor [1], the GSPA+QR gene embedding groups cell type-specific genes identified in PanglaoDB [31]. In a dataset of embryoid body (EB) cell lineages [69], one trajectory within the gene embedding reveals an association with the hemangioblast lineage identified in the original work, where genes early in the trajectory are associated with embryonic stem cells (*NANOG, POU5F1*), and hemangioblast-specific genes are later in the trajectory (*CD34, PECAM1, TAL1*).

#### Gene localization assessment from embeddings

Beyond gene-gene coexpression, we benchmarked embeddings for the ability to capture how differentially localized each gene is, where gene signals that are localized are considered more relevant for describing cell-cell variation, and gene signals that are more diffusely expressed are less relevant for describing cell-cell variation (Figure 3a). While simulated datasets consist of gene signals that vary in localization, we do not have ground truth scores for these genes. Therefore, for the gene localization task, we used the same simulated datasets to construct the cell-cell graph, then generated and embedded artificial graph signals of varying localization. We derived “ground truth” localization scores for these artificial signals based on the insight that we could constrain the size of the region where signals were defined, termed “window”. The size of this window is inversely related to the true localization score (Methods, Extended Data 8). Then, to predict signal localization, we embedded the uniform signal and computed the score as the multiscale distance to the uniform signal (as described in Definition 2), where the predicted localization should be highly correlated with the true localization score (Figure 3b). GSPA+QR outcompeted baselines for all three simulated datasets, followed by GSPA (Figure 3c, Extended Data Figure 9).

**Figure 3.**
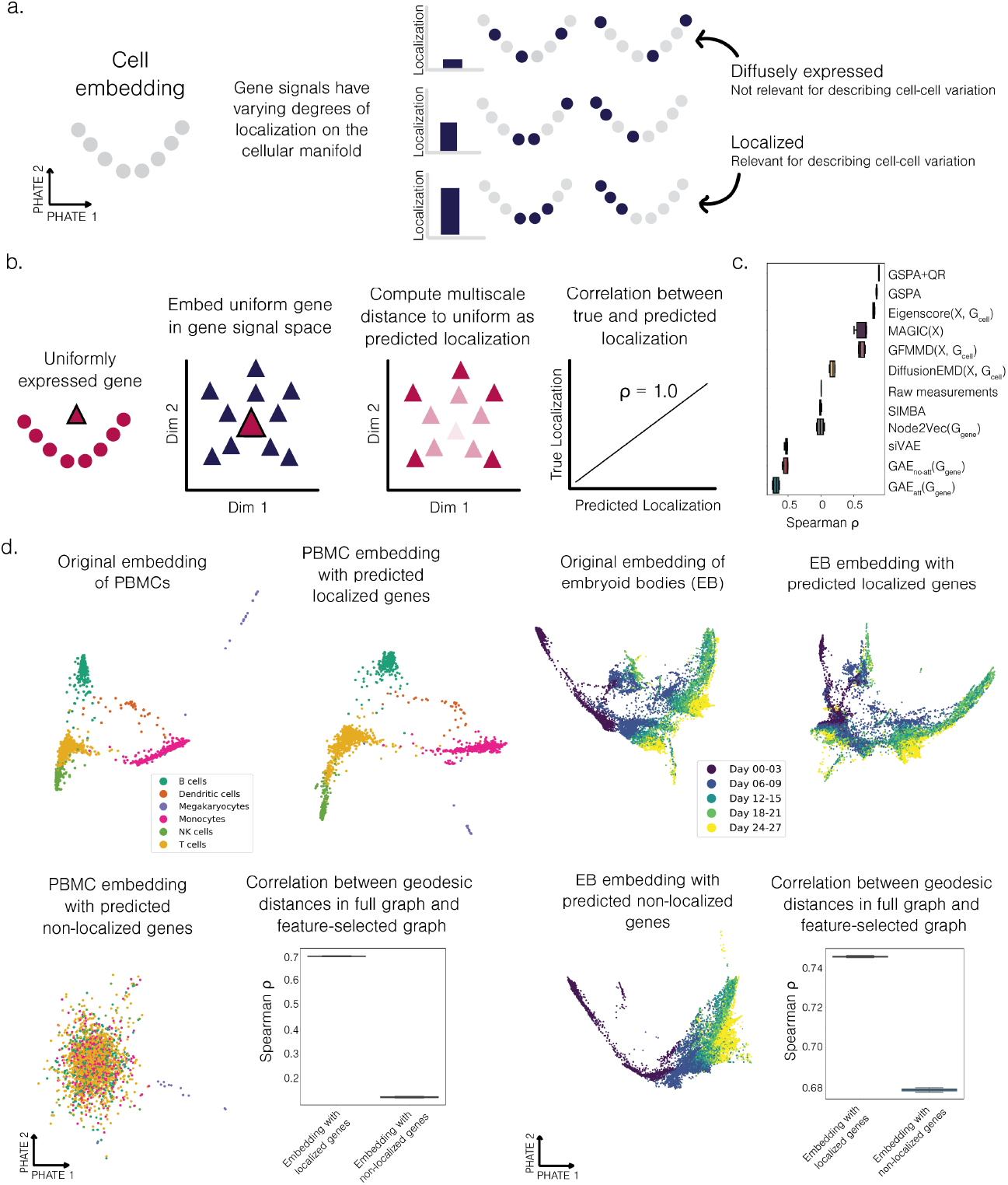
GSPA enables analysis of differential localization of signals. a. Differential localization diagram. b. Diagram of how localization reveals genes that are most distant from uniform. c. Spearman correlation evaluating performance for all comparisons across 3 runs. d. Original cell embedding versus cell embedding generated with predicted localized genes or predicted non-localized genes only; correlation between geodesic distances in original cell-cell graph versus feature selected cell-cell graphs. Shown for PBMCs (left) and EB data (right).

As localized genes are informative for describing cell-cell variation, we hypothesized that predicted localized genes can be used for topologically-informed feature selection (Methods). For the PBMC dataset, we visualized the cells first using all genes, then only the top predicted localized genes. This revealed a strikingly similar embedding of cells and separation of cell types, suggesting that the top localized genes are able to capture the majority of the cell type variation and preserve information about the entire transcriptional space of the cells. By contrast, visualizing the PBMCs using the least localized genes completely removes all cell type variation, as these genes likely do not describe meaningful differences between cells. The correlation between pairwise geodesic distances in the cell-cell graph derived from all genes versus selected genes reveals that localized genes are able to better preserve information about local and global relationships between cells than non-localized genes (Figure 3d).

Similarly, for the embryoid body dataset, the predicted localized genes are able to capture the major and minor trajectories within the embedding based all transcriptional measurements, whereas the predicted non-localized genes only capture the major axis of variation (the timecourse). Again, correlation between pairwise geodesic distance in the full cell-cell graph versus the feature-selected cell-cell graphs shows stronger preservation of cell-cell distances by localized genes (Figure 3d).

#### Robustness to preprocessing steps and graph construction

To evaluate robustness to steps to process the cellular measurements before mapping the gene space, we ran our coexpression experiment and localization experiment for all comparisons over each combination of the following: two runs, two single-cell dataset transformations (`log` and `sqrt`), four choices for *k* in construction of *k* nearest neighbors graph (5, 15, 25, 50), and three choices for construction of nearest neighbors graph (*k* nearest neighbors (*k*NN), shared nearest neighbors (SNN), and construction with an adaptive *α*-decaying kernel). Together, this resulted in 48 runs for each method (Methods, Extended Data Figure 10b i.). On average across all hyperparameters and preprocessing choices, GSPA and GSPA+QR again outperformed all other approaches (Extended Data Figure 10b ii.). Furthermore, despite potential sensitivity to graph construction, approaches that leveraged the cell-cell graph to calculate gene-gene relationships outranked approaches that used pointwise gene measurements on both experiments (Extended Data Figure 10c). For the coexpression experiments, approaches with the cell-cell graph had an average rank of 2.929, and approaches without the cell-cell graph had an average rank of 8.071 (Extended Data Table 1a). For the localization experiments, approaches with the cell-cell graph had an average rank of 2.686, and approaches without the cell-cell graph had an average rank of 8.314 (Extended Data Table 1b). This result reinforces the desired distance preservation and noise robustness properties garnered from using the cell-cell graph and further supports our assertion that considering genes as signals on the cell-cell graph can improve analysis of gene-gene relationships. Additionally, as most single-cell sequencing analysis tools and pipelines construct a cell-cell graph, including for visualization, clustering, and trajectory inference [13,101], using the same graph can ensure consistent biological analysis with GSPA.

### 2.6 Gene embeddings reveal coexpression relationships in CD8+ T cells

We first chose to study CD8+ T cell differentiation in response to infection. T cells adopt a range of differentiation states in acute and chronic viral infections, creating diverse and heterogeneous substates differing by phenotype, physiological role, morphology, proliferative potential, and location [20]. Such heterogeneity has been established as a key feature of T cell immunity, where cells can differentiate between states in a “plastic” way. However, the gene signaling pathways and relationships that characterize these transitions are not fully known and can provide new insights for therapeutic intervention.

We therefore investigated a newly developed dataset comprising 39,704 sorted CD8+ Tetramer+ T cells, sequenced at three timepoints (Day 4, Day 8, and Day 40) from two conditions (acute LCMV and chronic LCMV infections) (Methods). We preprocessed this data and visualized the cellular manifold with PHATE, revealing distinct structure that does not separate into clear clusters or trajectories, as expected due to the dynamic and multiscale nature of T cell heterogeneity. Clustering cells and identifying top differentially expressed genes (DEGs) shows overlapping genes across clusters and often genes that are highly expressed in all clusters, such as *Rps19* and *Rps20* (Extended Data Figure 11a, Extended Data Figure 11b). Visualizing known markers of T cell differentiation highlights that even relevant genes are not specific to a particular cluster, and some gene signatures, such as genes related to proliferation (e.g. *Mki67*), overlap others, such as genes related to activation (e.g. *Gzma*) (Figure 4a, Extended Data Figure 11b). This motivates mapping the gene space to capture key signatures across conditions and timepoints.

We computed representations for the top highly variable genes using GSPA+QR. We then visualized the gene embedding and clustered genes into six modules (Methods, Figure 4b, Extended Data Table 2). Next, we computed the localization score of each gene and visualized this score on the gene PHATE (Figure 4c). Genes that had higher localization scores were on the periphery of the embedding, which fits our intuition, as genes that are more spread out are likely to be less committed to one particular population or region of the cellular graph. Furthermore, we saw little relationship between the localization score and number of cells expressing each gene (Spearman *ρ* = −0.159) or the mean overall expression for each gene (Spearman *ρ* = −0.108), indicating that the score is capturing a meaningful structural property of the signal.

To better compare and contrast differential localization and differential gene expression in real-world data, we developed a cell clustering-based ranking of genes using the maximum z-score underlying the computation of the p-value for each gene across all clusters (Methods). This ranking, which reflects how enriched each gene is, is slightly positively correlated with the localization score (*ρ* = 0.327) (Figure 4d). Genes that had high cluster-based rank and high localization included genes strongly enriched in a particular cluster (e.g. *S1pr5*). Genes that had low cluster-based rank and low localization included genes with low or non-specific enrichment overall (e.g. *Trav13d-1*). Other genes, such as *Rps20*, were ranked highly by the cell-clustering rank but lowly by the localization rank due to high mean expression in one cluster but high overall expression and spread throughout the manifold. Finally, some genes, including *Tox*, were ranked highly by localization but lowly by cell-clustering rank due to their varied but specific enrichment across the T cell manifold. Together, this demonstrates the ability for localization to not only capture cluster-specific signatures without prior clustering but also prioritize more subtle, lowly-expressed signatures and deprioritize ubiquitously expressed genes. In addition, we show that re-embedding cells with the top localized genes better preserves the overall manifold structure than re-embedding cells with the bottom localized genes, highlighting localization for topologically-informed feature selection (Extended Data Figure 12).

Next, we characterized each gene module based on marker genes and gene set enrichment scores, which did not conform to distinct cluster-like shapes (Figure 4e). We identified cells with strong enrichment for each gene module and compared the proportion of these cells from each condition, which revealed many expected patterns from existing literature [25, 35]. Gene module 0 seemed to capture a memory-specific signature as it was most highly enriched in the Acute Day 40 timepoint and contained hallmark genes such as *Il7r* and *Bcl2* [37]. Gene module 1, enriched for naive and memory genes, had a higher proportion of cells from Acute Day 4 and Day 40 timepoints. Module 2, with proliferation genes, was enriched in Acute Day 4 and Chronic Day 4. Module 3, with effector genes, was enriched in Acute Day 8 and Chronic Day 8, and module 4, with late effector and exhaustion genes, was enriched in Acute Day 8, Chronic Day 8, and Chronic Day 40 (Figure 4e). Module 5 captured an interferon response signature, which was interestingly present in both Acute and Chronic Day 4, but only seemed to persist past that timepoint in the Chronic setting (Figure 4e). This is in line with findings that showed type 1 interferon can enforce T cell exhaustion in chronic infection settings [68, 86, 102].

While gene modules group genes based on relatively similar expression profiles, the localization score determines how specific that expression profile is. For example, *Rps20* and *Tcf7* both belong to gene module 1, but, because *Rps20* shows high expression in other cells, whereas *Tcf7* shows almost no expression in other cells, *Tcf7* has a higher localization score. Therefore, to further characterize each gene module, we selected the most highly localized genes and built a *k*-NN graph, highlighting the strong gene-gene coexpression relationships in each module (Extended Data Figure 11d). For example, the naive/memory gene-gene graph contained edges between *Tcf7, Ccr7, Sell, Gpr183*, and other known markers including those associated with acute and chronic progenitor cell states such as *Id3* [104] and *Xcl1* [23]. The effector module contained edges between *S1pr5, Zeb2, Gzma, Gzmk*, and others, and the exhaustion module contained edges between *Pdcd1, Cxcr6, Lag3, Tigit*, and others. Protein-protein interaction analysis with STRINGdb [88] revealed that all modules showed significantly higher interaction than expected for a random set of proteins of the same size and degree distribution (*p <* 1.0*e*-16 for all modules), suggesting each gene module represents a biological signature validated in prior literature.

To better understand if GSPA is facilitating biological discovery beyond existing approaches, we constructed an experiment based on gene set enrichment of type 1 interferon signaling (Methods). The top differentially expressed genes from each cell cluster show no enrichment of type 1 interferon signaling, despite its known relevance to T cells in chronic conditions (Figure 4f). Similarly, the top localized genes in the gene module of interest from cNMF and gene embedding approaches show low enrichment scores. By contrast, the top localized genes in the gene module of interest from GSPA and GSPA+QR reveal strong enrichment, suggesting gene module and localization analysis with GSPA can uniquely identify this relevant signature.

Finally, to further test the utility of gene-gene *k*-NN graphs built from the gene representations, we compared the gene-gene graphs for a negative control of Acute Day 8 cells versus cells from a *Tbx21* knockout (KO) and a *Klf2* KO experiment, visualizing only gene-gene interactions that were knocked out in the experiment but existed in the negative control (Figure 4g, Extended Data Figure 11e, Methods). This uncovered interesting gene networks lost in the *Klf2* KO such as those around *Bach2* and *Id2*, which have been implicated in memory and effector differentiation respectively [83, 104]. Interestingly, in the *Tbx21* KO network, *Cd69* and *Batf* appear, which may be reflective of *Tbx21* ‘s role in enforcing effector differentiation [48]. Such analysis highlights the ability of gene signal pattern analysis to capture gene coordination events in perturbation-specific ways.

### 2.7 GSPA captures cluster-independent patterns of cell-cell communication in tissue-resident immune cells

One of the key benefits of learning informative gene features is the ability to characterize patterns of gene signals beyond the boundaries of cell states, including identifying signatures within or between cell states. This is especially useful for characterizing patterns of cellular communication, as these interactions can involve multiple cell states or a small subset of cells that are responding strongly to a stimulus. In canonical cell-cell communication analysis, which calculates an enrichment score based on how specific an interaction is between two cell states, both of these types of interactions are not likely to be prioritized [5].

Using known ligand-receptor (LR) pairs, GSPA-LR first obtains ligand and receptor embeddings individually, then concatenates them into a LR pair representation. We do this because two communication patterns *a* and *b* should be represented similarly if ligand_a_ and ligand_b_ show similar expression profiles and receptor_a_ and receptor_b_ show similar expression profiles. So, mapping the concatenated representations identifies LR pairs with shared patterning on the cellular manifold across and within cell types and is more informative than aggregating the ligand and receptor signals (such as through summation or averaging the expression of both). If pathway attributes are available for each LR pair, we can further characterize pathway-pathway relationships by building a *k*-NN graph of the LR pair representations 𝒢_*LR*_ = (*V*_*LR*_, *E*_*LR*_) and, for each pathway, defining a set indicator signal **1**_*pathway*_ on all vertices of 𝒢_*LR*_, where **1**_*pathway*_(*v*) = 1 if *v* ∈ pathway and **1**_*pathway*_(*v*) = 0, otherwise for *v* ∈ *V*_*LR*_. Using GSPA, we map the pathway space (Figure 5a).

We tested GSPA-LR in a recently published work examining the role of PD-1 in skin tolerance and potential mechanisms of checkpoint inhibitor-driven immune-related adverse events (Methods, [24]). The data supported the hypothesis that PD-L1 expression by myeloid cells functions to maintain healthy tissue through interaction with local CD8+ T cells, and PD-1-blocking immunotherapies result in adverse events. Given this interaction was specific to antigen expression and functionally validated, we sought to recover the PD-1/PD-L1 interaction axis using our pipeline.

The scRNA-seq dataset from this work comprises 21,178 skin cells from three conditions: antigen off (NO AG), antigen-expressing (AG), and antigen-expressing treated with checkpoint inhibitors (AG CPI) (Figure 5b). This dataset was previously annotated with four cell types (Figure 5c). As this dataset showed distinct cell types, we calculated cell type association scores across the gene space for each cell type. The gene embedding showed distinct regions of the gene space enriched for distinct cell types (Extended Data Figure 13a). Further, the top genes most closely associated with each cell type were strongly and specifically enriched (Extended Data Figure 13b).

Despite separation of cell types on the transcriptional level, many key communication patterns showed enrichment within subsets of the cell types, or across more than one cell type. CCL5 is enriched in epithelial, myeloid, and T cells, and CCR5 is primarily enriched in myeloid cells and T cells. Both the ligand and the receptor are activated in response to the presence of antigen (AG, AG CPI). Additionally, PD-L1 is enriched in myeloid cells, and PD-1 is enriched in T cells, also activated in the presence of antigen (Figure 5d). We show that the commonly used cell-cell communication permutation test enabled by CellPhoneDB [29] captures significant interactions between CCL5 in myeloid cells and T cells to CCR5 in all cell types, but fails to reflect the CCL5 enrichment in epithelial cells and the condition-specific nature of the interaction. For PD-L1 and PD-1, CellPhoneDB captures the low expression of PD-L1 in all cell types, but it is unclear from these results alone that PD-L1 is strongly enriched in myeloid cells in the presence of antigen (as validated in [24]) (Figure 5e).

By contrast, GSPA-LR ignores cell type labels and reveals LR pair representations and modules capturing a diverse range of ligand and receptor profiles (Methods, Figure 5f, Extended Data Table 3, Extended Data Figure 14). Module 5, which includes CCL5-CCR5, CCL4-CCR5, IFNG-IFNGR1, and ITGB2-ICAM1, shows ligand enrichment in the AG and AG CPI subsets of the epithelial, myeloid, and T cells, and receptor enrichment in the AG and AG CPI subsets of the myeloid and T cells. Gene set enrichment analysis indicates enrichment for adhesion and diapedesis, as well as binding of chemokines to chemokine receptors. These processes are not cell-type specific, but rather condition-specific migration and effector activity in response to the skin lesion. Module 19, which includes PD-L1-PD-1, PD-L2-PD-1, CXCL16-CXCR6, and CCL12-CCR2, shows ligand enrichment in AG and AG CPI-specific myeloid cells, and receptor enrichment in AG and AG CPI-specific T cells, matching the known pattern described in the original work. Module 19 is enriched for integrin activity and PD-1 signaling, both associated with T cell activation and exhaustion (Figure 5g). Finally, GSPA-LR maps pathway relationships and reveals the colocalization of several related pathways, including those related to PD-L1/2 and T cell cytokine and chemokine secretion (Figure 5h). GSPA-LR was therefore able to capture both pair-pair and pathway-pathway relationships, demonstrating the ability to perform multiscale gene signaling analysis.

### 2.8 Spatially-colocalized gene modules correspond to key communication hubs

Gene signal pattern analysis can also derive feature embeddings for data modalities beyond scRNA-seq (GSPA-multimodal). Integrated diffusion [56] creates a joint diffusion operator **P**_*integrated*_ from multimodal data, or data gathered from different measurements on the same system. GSPA-multimodal uses this integrated diffusion operator to construct an integrated wavelet dictionary, outputting feature representations informed by all modalities (Methods, Figure 6a).

Here, we demonstrate the capability of GSPA-multimodal on spatial transcriptomic data, where both gene expression and spatial coordinates are measured for spots consisting of on average 10 cells. Spatial transcriptomics is a powerful tool for adding additional organizational context to gene expression analysis. Biological systems such as the lymph node contain many highly active microenvironments and diverse cell types corresponding to spatially distinct substructures and unique organization. However, like scRNA-seq data, most spatial technologies are accompanied by a large amount of technical noise and artifact [58]. This obfuscates the ability to assess gene patterns and relationships. Additionally, spatial transcriptomic analysis is often interested in identifying “spatially variable” genes, or genes that describe spatial variation [87].

GSPA-multimodal constructs gene embeddings informed by both expression and spatial affinity between spots, and, through the localization framework, identifies spatially variable genes without bespoke models for spatial transcriptomic data.

We thus leveraged GSPA-multimodal to analyze a spatial RNA-sequencing dataset of a healthy human lymph node from 10x Visium (Methods, Figure 6b) [2]. We clustered the gene embedding and computed the localization score (Methods, Figure 6c, Extended Data Table 4). Visualizing the mean expression and top localized genes of each gene module on the spatial coordinates highlight the spatial localization of each signature (Figure 6d).

Spatially-variable genes identified by SpatialDE [87] correspond to localized genes derived by our approach, where the localization score is significantly higher for spatially variable versus non-spatially variable genes (Methods). However, by using both the expression graph and the spatial graph to determine localization, GSPA-multimodal is further empowered to identify relevant biology. Two example genes that are considered localized but insignificant by SpatialDE, *CD34* and *MMRN2*, are enriched in the adventitia of the vasculature and have been previously implicated as progenitor cells that may give rise to other fibroblast subsets [36] (Figure 6e).

Notably, while there is a large variety of cell types, this data is not at single-cell resolution. Therefore, to aid in characterizing cell types enriched across the tissue, we leveraged an integrated atlas of human secondary lymphoid organs composed of 73,260 cells from [52] and used the cell2location [52] mapping to determine the cell types enriched for each gene module. This reveals enrichment of monocytes, macrophages, mast cells, endothelial cells, FDCs, and VSMCs for gene module 0, which spatially corresponds to the blood vessel; GC-committed B cells, T follicular helper cells and follicular dendritic cells for gene module 1, enriched in the germinal centers of the tissue; T cells for gene module 2; non-GC-specific B cells for gene module 3, enriched in the B follicule; B plasma cells for gene module 4; and B IFN cells for gene module 5 (Extended Data Figure 15).

As each spot contains cells from different cell types with a high degree of cell-cell interaction and organization, we hypothesized that we could identify gene signaling corresponding to communication events. We thus leveraged the OmniPathDB [95], a knowledge graph containing curated intercellular and intracellular gene-gene signaling interactions. For each gene module, we built a *k*-NN graph from the gene embeddings, and we pruned the graphs to include only edges that exist in the OmniPathDB network. As OmniPathDB edges are directional, this also allows us to impute directionality to the gene-gene graph. Then, we sought to identify the cell types involved in the gene-gene signaling events. We repeat the following for each gene module graph: for each directed edge (*gene*_*s*_, *gene*_*t*_), for all pairs of cell states (*celltype*_*a*_, *celltype*_*b*_), if *gene*_*s*_ is differentially expressed in *celltype*_*a*_, and *gene*_*t*_ is differentially expressed in *celltype*_*b*_, we add a directed edge from *celltype*_*a*_ to *celltype*_*b*_. For gene modules with multiple cell types, this revealed complex communication patterns (intercellular edges in blue, intracellular edges in red). Endothelial cells, VSMCs, macrophages, and monocytes showed strong signaling in the blood vessel. B cycling cells and other GC-committed B cells were communicating in the germinal center. Finally, T cells showed highly active communication in the T cell zone (Figure 6f). In sum, GSPA-multimodal enables spatial analysis of dynamic and active cell-cell communication characterizing the human lymph node. Importantly, the integrated diffusion operator can naturally extend to any multimodal data, including scRNA-seq and sc-ATAC-seq, or datasets with multiple relational structures describing the same underlying phenomenon. In this way, gene signal pattern analysis extends beyond single-cell analysis to represent multimodal features.

### 2.9 Multiscale data features map patient manifolds and predict outcomes

Finally, gene signal pattern analysis can be used to map manifolds of patient samples (GSPA-Pt). Intuitively, we hypothesized that constructing patient vectors based on GSPA gene embeddings could capture both the topology of the cell-cell graph and the coexpression between genes for improved prediction of patient response. Additionally, as the individual features of the patient vector correspond to genes, we could explore the genes most predictive of response and non-response.

To this end, in the GSPA-Pt framework, we first consider 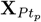 as a single-cell dataset for patient *p* for *p* ∈ 1…*P*. We then concatenate all samples to build a shared cell-cell graph 𝒢_*cell*_, which we use to build the wavelet dictionary 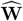 as before. As each entry in 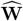 is associated with a patient *p* ∈ 1…*P*, we can split 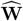 into patient-specific dictionaries 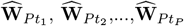. Then, for each *p*, we project 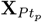 onto 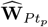 and learn a reduced patient-specific gene representation. Each patient is represented by a gene embedding, which is flattened into a vector for downstream analysis (Methods, Figure 7a).

We test GSPA-Pt in a dataset of 48 melanoma samples pre- and post-checkpoint immunotherapy [77]. Despite the successes of checkpoint immunotherapy in melanoma, many patients fail to show a response. The authors show that the immune cells in the tumor microenvironment play a key predictive role in distinguishing responders and non-responders. Therefore, we first sought to map each patient sample, corresponding to a scRNA-seq dataset of sorted and sequenced CD45+ cells. Then, we evaluate the classification of response based on this representation using a logistic regression classifier.

For comparison, we learn patient sample embeddings in four other ways. First, we build the combined cell-cell graph 𝒢_*cell*_ and consider the patients as set indicator signals on the cells derived from each patient sample. We then use GSPA to represent patient signals. Second, we compute the mean expression across all cells through aggregation for each patient. Third, as the cells were clustered and annotated in the original work, we compute the proportion of each cell type and define patient features based on these proportions, similar to [16]. Finally, we do the same for each cell subtype within CD8+ T cells, which were previously shown to be highly predictive of response.

Visualization of the patient embeddings shows that the GSPA-Pt gene embeddings, GSPA patient signals, and mean expression separate responders and non-responders, whereas the proportion of cell states and substates do not do this as distinctly (Figure 7b). Classifying response based on the GSPA-Pt gene embeddings showed the highest performance, followed by the mean expression embeddings (Figure 7c). Interestingly, though the gene-based embeddings outperformed the cell-based embeddings, GSPA on patient signals outperformed the cluster proportion representations, suggesting that learning multiscale features beyond clusters can improve response prediction.

Because the patient embeddings are comprised of gene features, the coefficients of the logistic regression classification reflect the importance of different genes for predicting response and non-response (Figure 7d, Extended Data Table 5). Many relevant genes were associated with T cell function, as expected due to the established role of T cells in recognition of tumor antigens and control of the tumor [74, 77]. Genes most associated with non-response include genes linked to T cell terminal differentiation (e.g. *Nkg7* (rank 1), *Gzma* (rank 5), *Cd38* (rank 28)), resembling programs identified in the original work [77], work on T cell exhaustion [33], and related work in the context of melanoma [90]. Genes most associated with response include genes linked to T cell progenitor states such as memory, activation, and survival (e.g. *Il7r* (rank 3), *Ccr7* (rank 4), and *Tcf7* (rank 16)). *Tcf7* +CD8+ frequency in tumor tissue was shown in the original work to predict response and better survival. These programs have also been associated with stem-like properties [79] and efficacy of diverse immunotherapies [59]. Together, these results reflect our understanding of the progenitor T cell state mediating the response and being the primary target of immunotherapy, as well as the inability of terminally differentiated cells to reacquire significant function [55, 82, 107]. By comparison, while the mean expression embeddings showed comparable gene rankings for some above markers (*Nkg7* (rank 3), *Il7r* (rank 17), *Ccr7* (rank 3)), it also ranked other important markers much lower, including *Gzma* (rank 115), *Cd38* (rank 176), and *Tcf7* (rank 499).

Building a patient manifold also provides information beyond T cell signatures. Five patient samples were visually separated from the remaining samples. Comparing the proportion of G1, the cluster annotated as majorly B cells, reveals that these patients’ cell samples are over 40% G1, far more than the average B cell proportion of the other patient samples (Figure 7e). Furthermore, patient manifolds allow us to analyze the overall shape of the patient space, which is useful for predicting or characterizing patient trajectories (Figure 7f). In this dataset, some patients had samples taken pre- and post-immunotherapy. Patient 1 was characterized as showing a clinical response of resistance [77]. Three samples were obtained from this patient: baseline (Day 0, lesion classified as responder (R)), post-therapy I biopsy (Day 48, R), and post-therapy II biopsy (Day 437, non-responder (NR)). Despite the baseline and post-therapy I biopsy classification as responder, the patient manifold shows that these samples embed near non-response samples and show a trajectory toward non-response, suggesting that the tumor microenvironment resembles non-responding and terminally differentiated cells. Indeed, only 54.8% of the CD8+ cells from the baseline biopsy were associated with memory and survival (in subclusters 4, 5, and 6 from [77]), and 37.1% in the post-therapy I biopsy. Three samples were also obtained from patient 3, who, through whole exome sequencing, was determined to have a *B2M* deficiency associated with resistance to checkpoint therapy in melanoma. *B2M* -deficient tumors are less susceptible to infiltration by cytotoxic T cells [76], so although all three samples were determined to be non-responders, the samples embed near responders on the patient manifold and reflect changes in proportions of memory and activated CD8+ T cells (baseline: 80.0%, post-therapy I: 68.4%, post-therapy II: 90.7%). Finally, two samples from patient 20, who showed mixed response to immunotherapy treatment, were determined as non-responsive. The patient manifold shows the trajectory from baseline to post-therapy, while still in the non-response region, is moving in the direction of response, suggesting the mixed response may be through intertumoral differences in immune infiltration [71] or the presence of both progenitor and terminally differentiated CD8+ T cells. These findings collectively underscore the utility of interpretable and predictive patient representations for the analysis of clinical outcomes.

## 3 Discussion

Though there has been much interest in characterizing gene-gene relationships, mapping the gene space has not been sufficiently investigated and motivated. In this work, we defined relevant comparisons from graph representation learning and graph signal processing, and we built experiments with three simulated datasets to evaluate gene embeddings, establishing the groundwork for future research in this space. We demonstrated the utility of framing genes as signals on the cell-cell graph, and we described our method of decomposing gene signals using diffusion wavelets, then reducing the dimensionality using an autoencoder framework. We also highlighted the superior ability of this technique to preserve gene-gene relationships, capture structure in the visualization, and reveal salient gene modules.

Additionally, we presented two new gene ranking approaches facilitated by this analysis. For datasets with clear cluster-like structure, cell type association rankings allow us to identify genes associated with each cell type. Alternatively, differential localization prioritizes genes that are “localized”, or specific to key regions on the cellular manifold, without any assumptions about its shape. Both of these rankings are broadly applicable to single-cell analysis, which primarily identifies important genes through statistical differences in mean expression between clusters. Usefully, by bridging localization and gene embeddings, we can identify the top genes most relevant to each gene signature separately. While there exist feature selection approaches for single-cell analysis [78, 98], it is not guaranteed that the top genes will display a diversity in gene trends or enrichment in different cell states. In our method, users can ensure diversity by choosing the top localized genes from each gene module.

We demonstrate the clear and broad utility of gene embeddings through the characterization of different datasets in several vignettes. In a scRNA-seq dataset of CD8+ T cells in acute and chronic infection at three timepoints, we reveal key gene modules that correspond to enrichment at different points of the T cell differentiation process. We also analyze gene-gene relationships through network analysis of a single-cell CRISPR screen, identifying coexpression events that were knocked out in the context of *Tbx21* and *Klf2*.

We additionally showed the utility of gene representations beyond characterizing the cellar state space. GSPA-based ligand-receptor analysis (GSPA-LR) demonstrated the ability to analyze ligand-receptor and pathway-pathway relationships without cell type annotation in a peripheral tolerance model [24], recovering condition-specific communication between subsets of multiple cell types. GSPA-multimodal on spatial transcriptomic data identified spatially enriched gene modules and microenvironmental signaling events within each module. Finally, GSPA-Pt mapped single-cell datasets from patients with melanoma, using the gene space to learn an interpretable representation and prediction of patient response.

This work opens up new avenues for further exploration. While we identified gene modules in Section 2.6 and Section 2.8, and ligand-receptor pair modules in Section 2.7, the embedding spaces showed continuous structure and may be further explored with gene trajectory analysis and gene archetypal analysis. One could also consider patterns of gene signals beyond localization; gene signals that are broadly pervasive and distributed throughout the cellular state space may be useful for identifying key marker genes for the system or housekeeping genes for future experiments. Calculating gene distance to other synthetic signals, such as cell pseudotime or a condition-specific signal, could reveal new insights into the particular biological system being explored. The gene signal pattern analysis architecture is also flexible and can be combined with supervised losses for jointly-trained gene embeddings to predict treatment response or cell state.

Importantly, this work represents a key contribution to graph feature or signal representation learning, and future work can leverage ideas presented here for a variety of graphs and signals. For example, localization scores can be used to characterize signals that are highly specific, which indicates specialization or commitment to a particular group of nodes in diverse settings. We thus expect numerous biological and non-biological data types to benefit from gene signal pattern analysis to extract rich feature representations from large-scale high-dimensional data.

## 4 Data availability

The accession codes to the data used for figure 4 will be provided before publication. scRNA-seq datasets for figure 5 are available from the Gene Expression Omnibus (GEO) database under accession number GSE228586. Visium data for figure 6 can be downloaded from the 10x website (via a function in the scanpy package). scRNA-seq data for figure 7 are accessible through accession number GSE120575.

**Figure 4.**
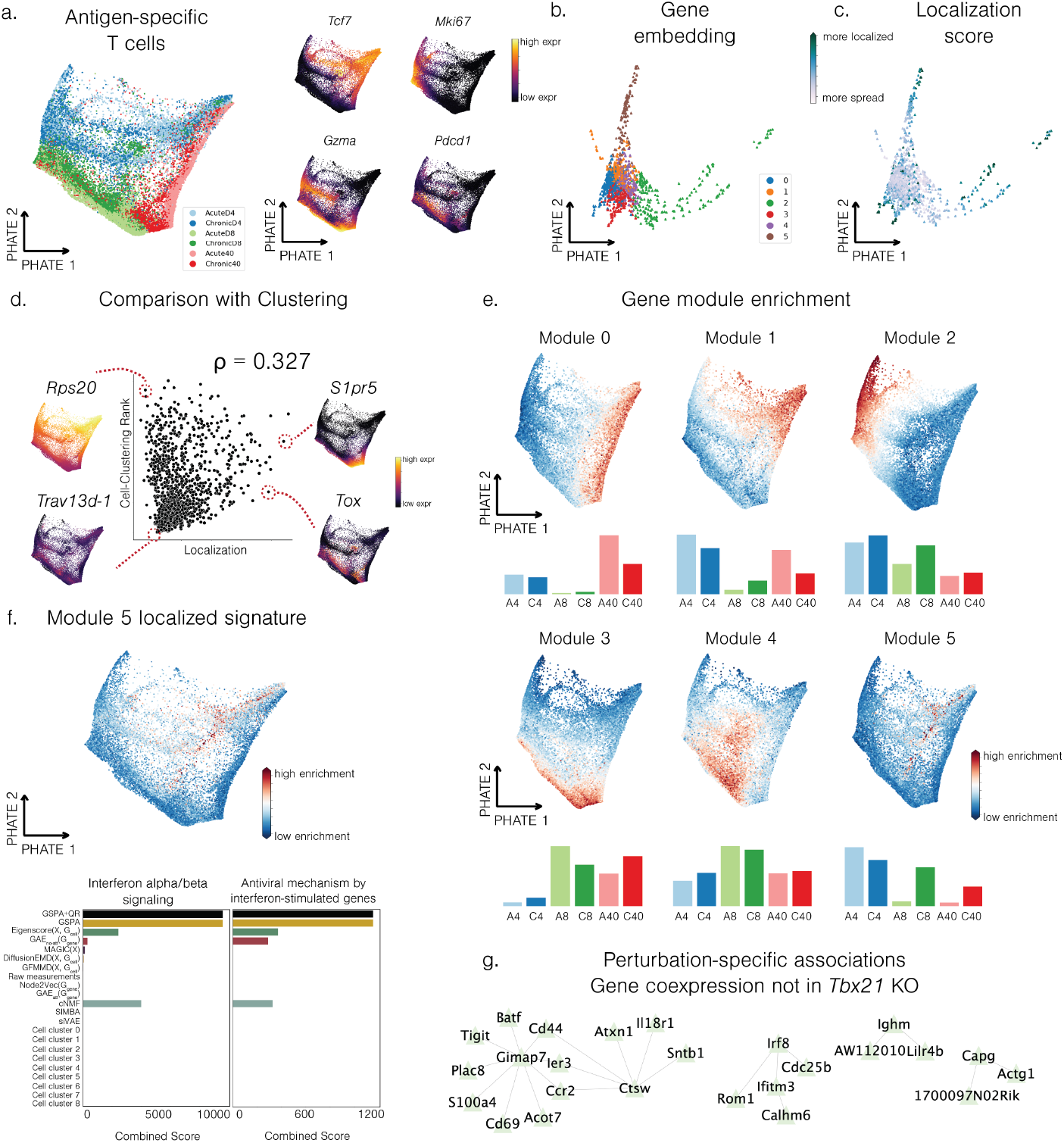
T cell gene signals reveal key signatures of differentiation. a. PHATE embedding of antigen-specific CD8+ T cells from six experimental conditions (left) and marker genes visualized (right). b. Gene embedding visualized with PHATE, colored by gene module assignment. c. Gene embedding visualized with PHATE, colored by computed localization score. d. Cell clustering rank versus localization score, with representative genes visualized to demonstrate similarities and differences. e. Gene module enrichment across all cells and per condition. f. Enrichment of top localized genes enriched in gene module 5 for GSPA+QR (top), and gene set enrichment scores for type 1 interferon gene sets for top genes from all comparisons (bottom). g. *k*-NN graph of gene-gene coexpression relationships that were knocked out in *Tbx21* knockout.

**Figure 5.**
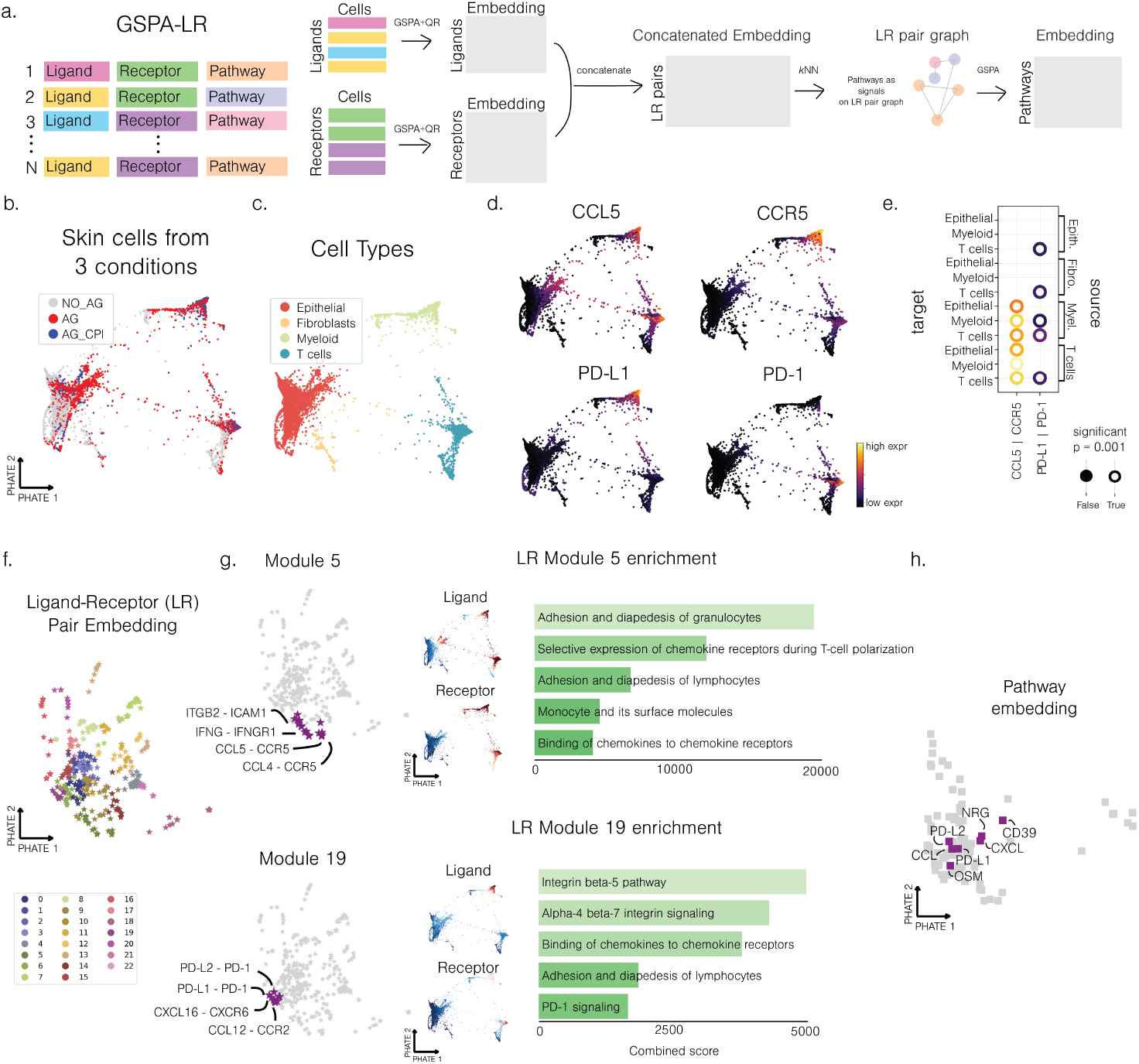
Identification of patterns of ligand-receptor communication in peripheral tolerance skin model. Schematic of GSPA-LR pipeline. b. Skin cells from no antigen (AG), antigen (AG), and antigen with checkpoint inhibitor (AG CPI) conditions visualized with PHATE. c. Skin cells colored by previously annotated cell types. d. Skin cells colored by CCL5, CCR5, PD-L1, and PD-1. e. Permutation test result from CellPhoneDB. f. Ligand-receptor pair embedding visualized with PHATE. g. Visualization of pairs, ligand and receptor enrichment, and gene set enrichment scores for module 5 (top) and module 19 (bottom). h. Pathway embedding visualized with PHATE.

**Figure 6.**
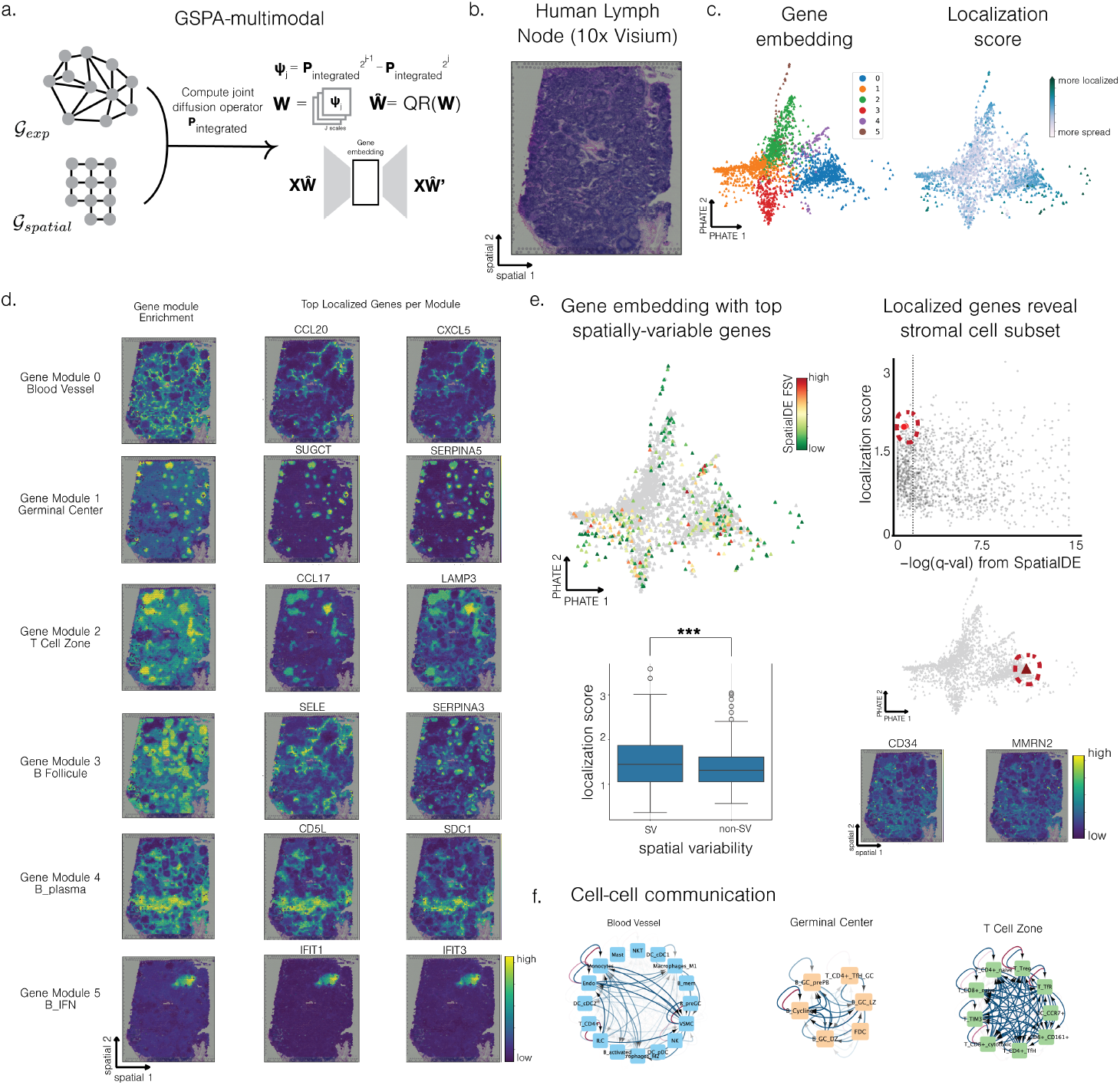
Gene embeddings capture communication pathways from 10x Visium human lymph node. a. Schematic of GSPA-multimodal using integrated diffusion on spatial transcriptomic data [56, 58] b. H&E stain of human lymph node tissue. c. PHATE visualization of gene embedding, colored by gene module assignment (left) and localization score (right). d. Enrichment of gene modules spatially and visualization of top localized genes. e. Gene embedding with top spatially-variable genes, where localization score corresponds with spatial variability (left). Localized genes that are not significant by SpatialDE reveal stromal subset (right). f. Cell-cell communication networks derived from gene-gene interactions with OmniPathDB.

**Figure 7.**
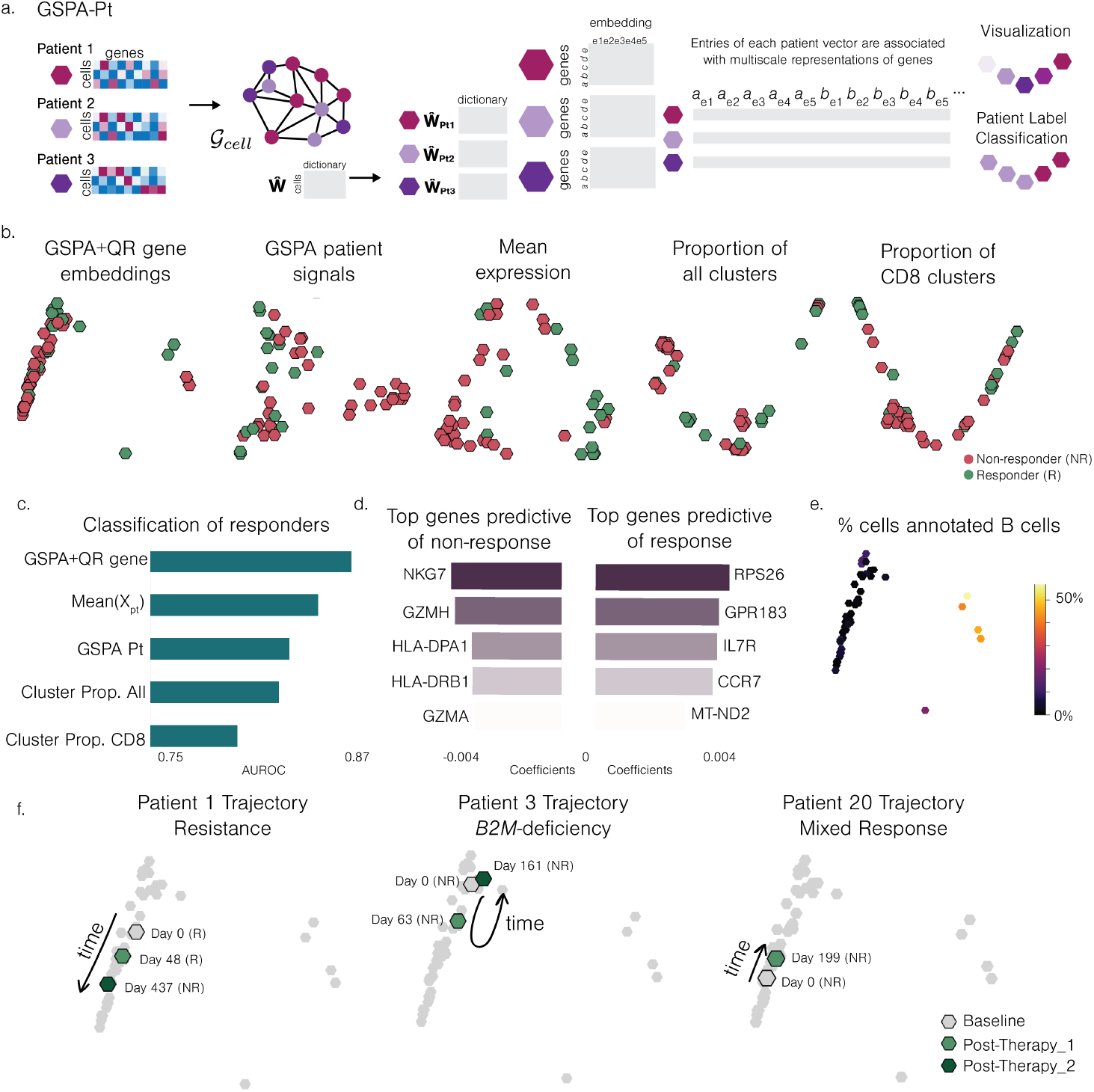
Patient manifold reveals response trajectories and biomarkers related to response prediction. a. Schematic of GSPA-Pt. b. PHATE visualization of patient embeddings based on GSPA+QR gene embeddings and comparisons. c. AUROC evaluation of response classification (logistic regression). d. Top genes predictive of response and non-response based on highest and lowest logistic regression coefficients. e. Patient embedding colored by percent of total cells annotated as B cells. f. Patient embedding with three samples from patient 1, 3, and 20 highlighted, corresponding to samples obtained from patients over time (pre-therapy baseline, post-therapy-1, and post-therapy-2). Trajectories of samples visualized.

## 5 Code availability

The source code for the Python package is available on The Python Package Index at https://pypi.org/project/gspa/ and on GitHub at https://github.com/KrishnaswamyLab/Gene-Signal-Pattern-Analysis. Notebooks to generate figures presented here are available at https://github.com/KrishnaswamyLab/GSPA-manuscript-analyses.

## 6 Methods

### 6.1 Related Work

#### Prior literature on examining gene-gene relationships

Effective strategies for identifying gene-gene relationships have been employed in bulk RNA sequencing data, including building gene coexpression graphs through gene-gene correlation [60, 61] or mapping from large databases [28]. However, gene-gene relationships resolved in bulk data are acutely sensitive to changes in sample cellular composition [30]. Single-cell transcriptomics avoids the compositional issues faced in bulk data, but constructing networks from these measurements directly is hindered by technical factors. Networks constructed without correcting for high levels of noise in gene expression measurements can be unreliable, as two genes that are related may not be expressed in the same cell. To overcome this problem, some studies first impute or denoise gene expression before calculating coexpression [14,27], which recovers relationships within a local neighborhood of the graph but does not account for both local and global distances between gene signals. Others have mapped genes to a high-dimensional space for improved prediction, which does not resolve low-dimensional embeddings for genes [7]. Finally, others filter predicted expression networks using additional evidence of interaction, such as the presence of a predefined transcription factor binding site among members of a co-regulated network [3].

#### Prior literature for patient sample comparison

Patient comparison of single-cell samples has historically compared cellular abundance [77]. Some approaches have built upon this idea to represent each patient as a vector of relative cluster proportions, which can then be directly compared and visualized [16, 57]. More recently, there have been methods that go beyond cluster abundances by modeling each patient dataset as a distribution on a common graph, enabling comparison via optimal transport between distributions [93, 105]. Patient embeddings can demonstrate how variation between patients corresponds to response to treatment. This becomes relevant for the development of predictive models of clinical outcomes, which can then be interpreted for the identification of predictive features [99].

### 6.2 Manifold learning and Diffusion Geometry

The manifold hypothesis is the belief that many high-dimensional datasets have a low-dimensional intrinsic structure. More specifically, given a dataset *X* = {*x*_1_, …, *x*_*n*_} ⊂ℝ^*m*^ of high dimensional observations, the manifold hypothesis assumes that the datapoints lie upon a *d*-dimensional manifold (i.e., a *d*-dimensional subset of ℝ^*m*^ that is locally equivalent to ℝ^*d*^) for some *d*≪*m*. For instance, in high-dimensional, single-cell experiments, each cell is described by a vector of counts per feature, where the number of features range from tens of key markers to thousands of genes. The measured cells make up the “cellular state space”, representing different possible cell states defined by the experimental setup. Redundancy in these measurements and high levels of coordination between the genes puts constraints on cellular behavior and reduces the effective degree. Thus, arbitrary points in ℝ^*m*^ do not correspond to plausible cellular states. Instead, the set of plausible states can be modeled as lying on a lower-dimensional manifold (see [43, 70, 99] for further discussion of modeling single-cell data on a manifold).

Many popular manifold learning methods are based upon constructing a graph 𝒢 = (*V, E*), whose vertices are the datapoints *x*_*i*_, which serves as a discrete approximation of the (unknown) underlying manifold. Weighted edges are constructed using a kernel *K* such as the Gaussian kernel

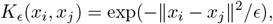

and one then defines a weighted adjacency matrix **A**∈ℝ ^*N×N*^ by *A*_*i,j*_ = *K*_*ϵ*_(*x*_*i*_, *x*_*j*_). The parameter *ϵ*, sometimes referred to as the bandwidth, controls the scale of the kernel, based on the idea that Euclidean distances ∥*x*_*i*_− *x*_*j*_∥ are equivalent to manifold distances (lengths of shortest path along the manifold) at small scales, but not at larger scales. Thus, *ϵ* should be chosen to be sufficiently small so that kernel only gives significant weight to intrinsically meaningful connections, but not so small that the graph becomes disconnected.

Given **A**, one can then define the diffusion operator **P** = **AD**^−1^, where **D** is the diagonal degree matrix (**D**_*i,i*_ = Σ _*j*_ **A**_*i,j*_, **D**_*i,j*_ = 0 if *i* ≠ *j*). This matrix **P** describes the transition probabilities of a lazy random walker exploring the vertices of the graph one step at a time. It is can also be interpreted as an operator to diffuse the values of signal **x** based on the values at neighboring points. Using higher powers of **P**, i.e. **P**^*t*^**x**, can be seen as averaging **x** over *t*-step random walks.

Based on these observations, Coifman and Lafon introduced diffusion maps [21], a method for embedding the datapoints into a low-dimensional Euclidean space ℝ^*d*^, *d*≪*m*, parameterized by a diffusion-time parameter which controls the scale (level of locality) of the representation. Formally, we take the eigenvalues 1 = *λ*_1_ ≥ *λ*_2_ ≥ · · · ≥ *λ*_*N*_ and (corresponding) eigenvectors 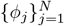 of **P** and map each point *x*_*i*_ ∈ *X* to an *d*-dimensional vector

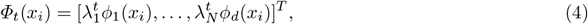

where *t* represents a diffusion time (note that the 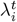 are the eigenvalues of the powered diffusion operator **P**^*t*^.) Small choices of *t* lead to more localized, small-scale representations, whereas larger values of *t* lead to larger-scale more global representations. However, in the latter case, we emphasize that these representations are larger-scale in the sense that they consider long-range dependencies between points *on the manifold* as measured by lengths of shortest paths, which are not the same as Euclidean distances.

Given the representations 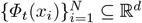, one may then analyze the data in reduced-dimension space, rather than the original high-dimensional ambient space. Indeed, using diffusion maps and related methods to model single-cell data on a manifold provides insight into key geometric and topological features of the cell-cell affinity graph, enabling characterization of the cellular state space despite sparsity, artifacts, and complex nonlinear relationships. As such, it has become a key tool in single-cell analysis for cell type identification [63], imputation [27], visualization [6, 69], differential abundance analysis [12], pseudotime calculation [8, 41], and multiscale analysis [57]. Below we will highlight two methods based on the diffusion maps style approach, MAGIC and Diffusion Wavelets.

MAGIC [27] is a denoising method based on the above mentioned diffusion operator. It is based on the intuition the many signals measured on, e.g., cell-cell graphs have extremely high noise levels, but that the noise typically exhibits high-frequency behavior. It is known that the 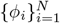 form a basis for ℝ^*N*^ and therefore any arbitrary graph signal **x** can be decomposed by

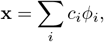

where the coefficient *c*_*i*_ represents how much of **x** lies at frequency *λ*_*i*_. One may verify that applying the diffusion operator yields

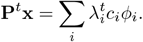

Additionally, we note that the eigenvectors are ordered in terms of smoothness/frequency with *ϕ*_1_ being the most smooth and each *ϕ*_*k*_ being smoother than *ϕ*_*k*+1_. Here, smoothness is quantified by

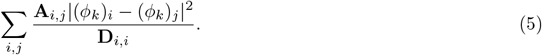

so that *ϕ*_*k*_ is smooth if (*ϕ*_*k*_)_*i*_≈(*ϕ*_*k*_)_*j*_ whenever there is a large edge-weight **A**_*i,j*_. In analogy to traditional Fourier analysis, we refer to the smooth eigenvectors, which vary slowly over the graph, as “low-frequency” and refer to the eigenvectors that vary rapidly over the graph as high-frequency.

Recalling that 1 = *λ*_1_ *> λ*_2_ *> λ*_3_ …, we see that **P**^*t*^ can be interpreted as a “low-pass filter” which preserves the low-frequency portion of **x** (corresponding to the larger eigenvalues, i.e., smaller values of *k*) and suppressing the higher frequency portion of the signal, which is typically noisy. This is the basis for the the MAGIC algorithm, which replaces the initial data matrix *X* (whose columns are the datapoints *x*_*i*_) with a denoised data matrix defined by

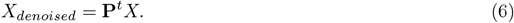

Diffusion Wavelets [22] aim to build upon the diffusion maps approach in order to produce a multiscale data representation. The diffusion map, defined as in (4), is based on the eigendecomposition of the powered diffusion operator **P**^*t*^ (since 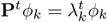). The diffusion time parameter, *t*, can be thought of as representing the scale of the representation since **P**^*t*^ performs a *t*-step random walk over the graph. In order to achieve a *multiscale* representation of the data, one can leverage multiple different values of *t*. This, along with classical wavelet constructions for data such as images (see, e.g., [66]), inspired Coifman and Maggioni to introduce diffusion wavelets of the form

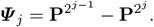

Since, the diffusion time parameter *t* represents the data scale, these ***Ψ*** _*j*_ model changes in the data across multiple scales and thereby produce a multiscale data representation. Since their introduction, these wavelets have played a powerful role in graph signal processing [10,72,81] and network anomaly detection [89], and have also been used to construct neural networks for graphs, manifolds, and other forms of geometric data [17, 18, 34, 75]. Additionally, we note that the choice of dyadic scales 2^*j*^ is was initially inherited from traditional Euclidean wavelets. However, [92], showed that this choice was unnecessary, and that the dyadic scales could be replaced with any increasing sequence of diffusion scales.

### 6.3 Gene signal pattern analysis (GSPA) detailed overview

GSPA can be described by the following steps explained in detail below:

1. Constructing a cell-cell similarity graph from single-cell data
2. Building a dictionary of graph diffusion wavelets from the cell-cell similarity graph
3. Projecting gene signals onto the wavelet dictionary
4. Learning a low-dimensional representation of gene signal projections with an autoencoder

#### Constructing a cell-cell similarity graph from single-cell data

To build the graph 𝒢_*cell*_ = (*V*_*cell*_, *E*_*cell*_), we compute the Euclidean distances between all pairs of cells and apply an *α*-decaying kernel to calculate affinities [12, 69]. The *α*-decaying kernel is defined as

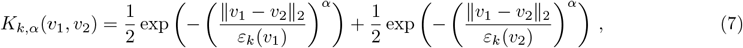

where *v*_1_ and *v*_2_ are cells ∈ *V*_*cell*_, viewed as points in ℝ^*m*^ corresponding to columns of **X**, *ε*_*k*_(*v*_1_), *ε*_*k*_(*v*_2_) are the distance from *v*_1_, *v*_2_ to their *k*-th nearest neighbors, respectively, and *α* is a parameter that controls the decay rate (i.e., heaviness of the tails) of the kernel. This construction generalizes the popular Gaussian kernel typically used in manifold learning while addressing some of its key limitations, as explained in [70]. This defines a fully connected and weighted graph between cells such that the connection between cells *v*_1_ and *v*_2_ is given by *K*(*v*_1_, *v*_2_). To increase computational efficiency, we sparsify the graph by setting very small edge weights (i.e., ≤1e-4) to 0. Additionally, **A** is defined as the adjacency / affinity matrix of 𝒢_*cell*_, or binarized *K*, and **D** is the diagonal degree matrix of 𝒢_*cell*_.

#### Building dictionary of graph diffusion wavelets for gene representation

Like many other tools from signal processing, wavelet analysis can naturally be extended to graphs and manifolds. In classical signal processing, for e.g., the analysis of temporal data, a wavelet dictionary is defined by taking a function *ψ* and a set of transformations of this function by time and scale. To adapt these methods to graphs, where there is no concept of time or linear space, we center wavelets at vertices on the graph and change scales via diffusion. Given the diffusion operator **P** (renormalized to have largest eigenvalue 1), we follow the construction of diffusion wavelets in [22]. Using ideas related to [9, 38], we use **P** to induce a multiresolution analysis, interpreting powers of **P** as dilation operators acting on functions, and constructing downsampling operators to efficiently represent the graph at fine-grained and coarse-grained resolutions.

Given the cellular graph 𝒢_*cell*_, we define **P** = **AD**^−1^ as the diffusion operator. Each wavelet of scale *j* centered at vertex *v* can thus be calculated by computing the matrix 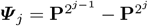 for 1 ≤ *j* ≤ *J* (and ***Ψ*** _0_ = **I** − **P**) and then extracting the *v*-th row via 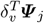, where *δ*_*v*_ is the Kronecker delta centered at the *v*-th vertex. Then 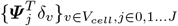 defines our wavelet dictionary **W** (where we use 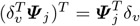 because our projection **X** →**XW** is performed via multiplication on the right). **W** is an *n* × *Jn* matrix (every wavelet takes values on the whole vertex set, and we have a wavelet for every vertex at each scale).

The number of scales for the wavelet dictionary *J* is defined as the *log* of the number of cells *n* based on the following lemma introduced by Tong et al. [93] and proven in the original work:

##### Lemma 1.

*There exists a K* = *O*(*log*|*V* |) *such that* 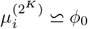 *for every i* = 1 …, *n, where ϕ*_0_ *is the trivial eigenvector of* **P** *associated with the eigenvalue λ*_0_ = 1.

This is based on the reasoning that if the Markov process converges in polynomial time with respect the number of nodes | *V*|, then one can ensure that beyond *O*(*log* | *V*|), all density estimates would be indistinguishable from each other.

Because the diffusion operator **P** is smoothing, we make the assumption that the numerical rank decreases as we take powers of the operator [22]. The faster the spectrum of **P** decays, the smaller the numerical rank of the powers of **P**, the more these can be compressed, and the faster the construction of ***Ψ*** _*j*_ for large *j*. Therefore, to decrease the size of the wavelet dictionary, remove redundant wavelets, and increase their interpretability, we can perform QR factorization. This results in a compressed wavelet dictionary 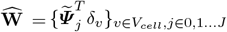, where for each *j*, 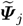 is a set of linear combinations of wavelets at *j* that account for the most variance. For large *j*, QR factorization naturally computes the numerical rank of 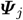 by taking a linear combination to form 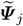 such that the total error in projecting 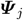 onto 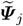 is less than some *ϵ* fraction of the norm of the whole 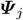. That is,

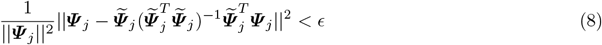

As raising **P** to the power of *t* diminishes the higher frequency-eigenvalues, diffusion wavelets support noise robustness and continuity properties by ensuring that 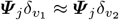 if vertex *v*_1_ is near vertex *v*_2_ on the graph. We test gene signal embeddings both with (GSPA+QR) and without (GSPA) QR factorization.

#### Projecting gene signals onto the wavelet dictionary

Given a gene signal **X**_*i*_ and wavelet dictionary 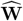 (or **W**), we project **X**_*i*_ onto the dictionary, which, for all gene signals, corresponds to 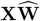 Alternatively, the solution can be framed as a set of *m* inner products provided by 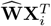 or 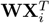 as a transformation of **X**_*i*_, or 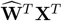 or **W**^*T*^ **X**^*T*^ for all gene signals. This transformation reveals each gene signal’s spatial and frequency information over the corresponding cell-cell graph 𝒢_*cell*_.

#### Learning a low-dimensional representation of gene signal projections with an autoencoder

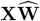is unnecessarily high-dimensional, where each feature corresponds to the gene signal projection of a particular cell at a particular diffusion scale. To reduce redundancy and improve computational tractability for downstream analysis, we reduce the dimensionality with autoencoder *D*○*E* where the objective is to minimize the mean squared error. That is,

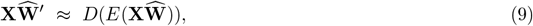

so that 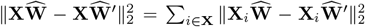 is as small as possible. The latent representation 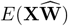 is the embedding we evaluate and characterize in downstream analysis, i.e.

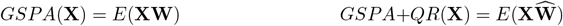

where **W** and 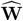 are uncompressed and compressed wavelet dictionaries (respectively), *E* is the encoder discussed above, and *GSPA* and *GSPA* + *QR* are taken to be the map *Θ* discussed in the problem setup.

#### Wavelet Projection and the Wasserstein distance

Earth Mover’s Distances (EMDs), alternatively referred to as Monge-Kantorovich or Wasserstein Distances, are a useful way of computing the distances between two signals. In the case where the signals correspond to probability distributions *µ* and ν, these distances can be thought of as the “cost” of moving a collection of points distributed according to *µ* to a collection of points distributed according to ν, where the cost of moving each point depends on the distance it must travel (defined with respect to some ground distance). In [91] (see also [93]), it was shown that the Wasserstein distance (with a truncated geodesic distance as ground distance) could be approximated by the Unbalanced Diffusion Earth Mover’s Distance (UDEMD) defined below. Here, we will show that the metric induced by our wavelets is continuous with respect to this UDEMD, i.e.,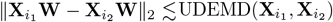.

In [91], the UDEMD [91] between two signals (genes) 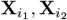 is defined as

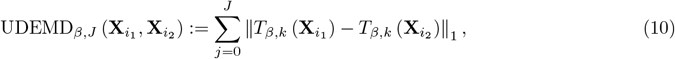

where 0 *< β <* 1*/*2 is a meta-parameter used to balance long- and short-range distances and *J* is the maximum scale considered here, and *T*_*β,j*_ is defined by

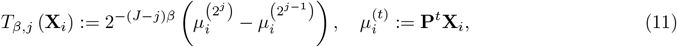

for 1≤ *j*≤ *J* and *T*_*β*,0_(**X**_*i*_) = 2^−*Jβ*^(**P**− **I**)**X**_*i*_.

The following result is a more detailed version of Theorem 1 first introduced in Section 2.3.

##### Theorem 1.

*For* 0 *< β <* 1*/*2, *the diffusion wavelet transform* **W** *(with maximal scale J) is Lipschitz continuous with respect to UDEMD*_*β,J*_, *that is there exists a constant C >* 0 *(depending on β and J and the ratio between the largest and smallest vertex degrees) such that*

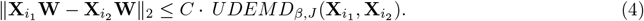

*Proof*. Let 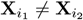. (The inequality (1) holds trivially in the case where 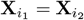.) We may compute

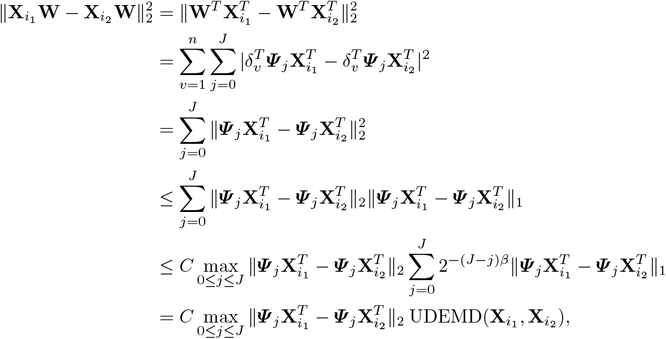

where *C* is a constant depending on *J* and *β*. It follows from Proposition 2.2 of [75] that

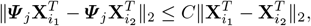

where *C* is a constant depending only on the ratio between the maximal vertex degree and minimal vertex degree. ([75] considers the wavelets on a weighted inner product space where vertices are weighted by degree. Transferring this result to the unweighted *ℓ*^2^ space induces dependence on the ratio between the maximal and minimal degrees.) Therefore, we have

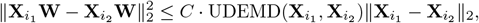

which in turn implies

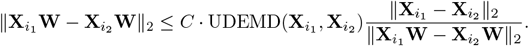

The lower bound in Proposition 2.2 of [75] implies that 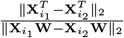 is bounded above by a constant (depending on the ratio between the maximal and minimal vertex degrees). Therefore, we have

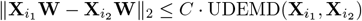

as desired.

### 6.4 GSPA for multiple modalities, datasets, and large graphs

#### GSPA for multiple modalities (GSPA-multimodal)

The approach described in Section 6.3 to construct cell-cell similarity graphs is useful for graphs derived from a single single-cell RNA-sequencing dataset. However, in cases where we have datasets of the same datapoints with multiple modalities, we can construct a combined representation using integrated diffusion [56]. GSPA-multimodal accepts two or more modalities and constructs affinity graphs for each modality (e.g., for two modalities, 𝒢_1_ and 𝒢_2_). Then, each graph has associated diffusion filters 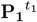 and 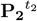, where *t*_1_ may not equal *t*_2_ due to differing degrees of noise, and are thus calculated by spectral entropy [56]. Finally, the integrated diffusion operator is calculated by multiplying diffusion filters, i.e. 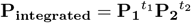. The integrated diffusion operator allows us to construct an integrated wavelet dictionary and project gene expression signals onto this dictionary for downstream analysis. Through the ability to flexibly define cell-cell affinity, GSPA-multimodal enables analysis of multimodal datasets including and beyond scRNA-seq.

#### GSPA for multiple datasets

For datasets consisting of cells from multiple datasets, GSPA can be used straightforwardly by concatenating all the datasets and constructing a graph of all the cells together (e.g. as done in Section 2.9). However, due to cells being sequenced in multiple runs, single-cell analysis is often affected by batch effects, where gene expression systematically differs between batches and confounds analysis of cell-cell variation. GSPA is also affected by batch effect if it is not corrected, where genes will separate based on association with batch in a way that confounds true gene-gene similarity. We show this with a simulated dataset generated with three (ground truth) clusters and two batches with batch effect (Extended Data Figure 1a). Gene embeddings learned with GSPA+QR show clear separation of genes enriched in each cluster through ground truth DE factor scores per cluster. However, these gene embeddings show clear separation of genes associated with each batch as well through coloring of ground truth batch effect factor scores per batch.

There exist a large number of batch correction methods, generally falling into three categories: integration in the gene space (where the output is batch-corrected gene expression measurements), the embedding space (where the output is batch-corrected cell embedding dimensions), or between graphs (where the output is a batch-corrected combined cell-cell graph) [64]. For batch correction in the gene space or the embedding space, the integrated output can be used to construct a cell-cell affinity graph and perform GSPA as above. For batch correction between graphs, the integrated graph can be used by GSPA, where we generate a wavelet dictionary based on the integrated graph. For example, we can use a mutual nearest neighbors (MNN) approach, based on the success of MNN-based correction introduced in [42], to construct a batch-corrected graph. In our simulated example, the integrated graph clearly shows no more separation of batches (Extended Data Figure 1b). GSPA+QR with this integrated graph still shows separation of genes based on enrichment per cluster, but now no more separation based on batch. We have implemented an extension of MNN-based correction to the adaptive decay kernel [12] to perform batch correction directly with GSPA.

#### GSPA for large graphs

For large graphs, Gene Signal Pattern Analysis utilizes diffusion condensation, a coarse-graining process which iteratively condensing datapoints toward local centers of gravity and is shown to approximate heat diffusion over the time-varying manifold [11]. Over the condensation time, the original coordinate functions are smoothed by a cascade of diffusion operators, which adaptively removes high-frequency variations. At each iteration, points closer than a given threshold ζ collapse to the same barycenter. This technique allows GSPA to summarize the underlying topology of the data manifold. Theoretical analysis of diffusion condensation from a topological perspective shows that the technique generalizes to, and is in some cases equivalent to, centroid-based hierarchical clustering [46]. However, this approach can provide more sensible clusterings due to the ability to coarse-grain without building a binary tree of the data, which can be arbitrary if many distances are equal. For GSPA, we use a version of diffusion condensation designed for single-cell analysis, Multiscale PHATE [57], which uses the potential representation of datapoints from PHATE [69] as the initial features.

For graphs larger than threshold *n*_*condense*_, we use Multiscale PHATE to iteratively condense datapoints to a small number of nodes. GSPA then filters for iterations with *n*_*condense*_ or fewer nodes, where each node represents a condensation of one or more cells. Finally, GSPA selects the iteration with a node count closest to *n*_*condense*_ to balance coarse- and fine-grained information. This represents a smaller cell-cell graph representing the same underlying manifold as the initial (larger) dataset, and GSPA computes a wavelet dictionary based on this graph. Then, gene signals are defined on the nodes of the condensed graph as the mean expression of all the cells in each node. By default, *n*_*condense*_ = 10, 000 cells. Due to the smaller size of the graph, computation becomes much more tractable with comparable results, where pairwise distances between genes from exact versus GSPA showed high correlation (R=0.900 for GSPA and R=0.713 for GSPA+QR) (Extended Data Figure 11c). With this approach, we can perform GSPA and GSPA+QR on 100,000 cells in 33.17 minutes and 30.18 minutes, respectively (Extended Data Figure 10a).

### 6.5 Computation of cell type association

For datasets where cells organize into distinct clusters, or cell types, gene signal pattern analysis can rank genes based on their specificity to each cell type. Given a dataset where each cell is assigned a cluster (or annotated cluster, i.e. cell type), for each cell type *C*, we can define a set indicator signal **1**_*C*_ on all vertices of the cell-cell graph, where **1**_*C*_(*v*) = 1 if *v* ∈*C* and **1**_*C*_(*v*) = 0, otherwise. Then, we can rank genes **X**_*i*_ based on how close they are to the normalized indicator signal 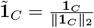 in their dictionary representation, i.e., how close 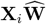 is to 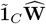. Formally, we define cell type association ranking of genes by the following score:

#### Definition 3.

*Given normalized cell type indicator signal* **1**_*C*_ *for cell type C and wavelet representation* **W**, *the cell type association score, c*(*i*) *for each gene signal* **X**_*i*_ *is defined as:*

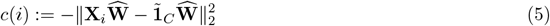

### 6.6 Computation of differential localization

Characterizing differentially expressed genes between clusters is not feasible for many biological systems. For example, for datasets that have trajectory-like structure, consist of subtypes within cell types, or do not organize into discrete populations, there is utility in identifying genes localized to particular areas of the cellular manifold without prior cell type identification. To this end, we naturally extend GSPA to a framework called *differential localization*. We calculate the specificity, termed gene localization score *l*(*i*), of a given gene signal *i* by calculating the multiscale representation of a uniform signal **u** and computing the distance between this and each gene signal representation. Genes are then ranked, where those that are most differentially localized are farthest from the uniform signal representation.

The gene localization score, *l*(*i*) for each gene **X**_*i*_, with normalized uniform signal 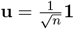 and wavelet Representation 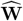, is defined as:

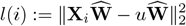

### 6.7 Comparison to other gene mapping strategies

Here, we describe in detail each comparison for our experiments with gene signal pattern analysis:

- Raw measurements approach embeds **X**.
- GAE_no-att_(𝒢_*gene*_) embeds 𝒢_*gene*_.
- GAE_att_(𝒢_*gene*_) embeds 𝒢_*gene*_.
- Node2Vec(𝒢_*gene*_) embeds 𝒢_*gene*_.
- MAGIC(**X**) embeds **X** after denoising with 𝒢_*cell*_.
- DiffusionEMD(**X**, 𝒢_*cell*_) embeds **X** via optimal transport on G_*cell*_.
- GFMMD(**X**, 𝒢_*cell*_) embeds **X** via MMD on G_*cell*_.
- Eigenscore(**X**, 𝒢_*cell*_) embeds **X** via alignment to Laplacian eigenvectors of _*cell*_.
- SIMBA co-embeds **X** and **X**^*T*^ via heterogeneous graph embedding.
- siVAE co-embeds **X** and **X**^*T*^ via jointly trained cell-wise and feature-wise VAEs.

We summarize and diagram these comparisons in Extended Data Figure 2.

#### Direct embedding of gene expression measurements

The simplest and most intuitive approach to map the gene space is with the original measurements. **X** consists of values where each cell is measured as a vector of gene expression counts, so we can consider the case where the genes are observations, and each gene is measured as a vector of expression counts in each cell. We use autoencoder *D* ○ *E* to reduce the dimensionality, where **X** ≈ *D*(*E*(**X**)) and *E*(**X**) is the embedding.

#### Embedding constructed gene-gene graph

Another approach is to construct a gene-gene *k*-NN graph 𝒢_*gene*_ = (*V*_*gene*_, *E*_*gene*_) from **X**, where each node in *V*_*gene*_ corresponds to a gene and each edge *E*_*ij*_ in *E* describes the similarity between gene *i* and gene *j* based on Euclidean distance. We can then leverage graph representation learning to propagate information between gene-gene relationships and learn node embeddings. We test one shallow embedding Node2Vec(𝒢_*gene*_), and two graph autoencoder embeddings. The graph autoencoder *D*_*no*−*att*_ ○ *E*_*no*−*att*_ consists of graph convolutional layers, where 𝒢_*gene*_ ≈ *D*_*no*−*att*_(*E*_*no*−*att*_(𝒢_*gene*_)). The graph autoencoder *D*_*att*_ ○ *E*_*att*_ consists of graph attention layers, where 𝒢_*gene*_ ≈ *D*_*att*_(*E*_*att*_(𝒢_*gene*_)). *E*_*no*−*att*_(𝒢_*gene*_) and *E*_*att*_(𝒢_*gene*_) correspond to the embeddings without and with attention, respectively.

#### Imputing gene signals with cell-cell graph

The above methods do not use information from the cell-cell graph for the computation of gene representations. Based on our desired properties (Section 2.1), we hypothesized that incorporating cellular affinities would enable the comparison of non-overlapping gene signals across local and global distances on the cellular manifold.

First, we compare against MAGIC [27], which imputes missing gene expression via data diffusion. MAGIC calculates a diffusion operator **M** powered to *t*, and left-multiplies **M**^*t*^ to **X**^*T*^ as a low-pass filter. For comparison, we left-multiply **X** to **M**^*t*^, which practically denoises gene signals and performs comparatively to MAGIC (data not shown). We then employ an autoencoder *D E*, where **XM**^*t*^ ≈ *D*(*E*(**XM**^*t*^)) and *E*(**XM**^*t*^) is the embedding.

#### Optimal transport distances between gene signals

Due to the relationship between GSPA and Wasserstein distance (i.e. optimal transport), we compare GSPA against three approaches for fast optimal transport that have been developed and used for gene signals on the cellular graph.

Diffusion Earth Mover’s Distance (DiffusionEMD) [93] computes optimal transport based on multi-scale diffusion kernels. This construction is related to UDEMD [91] described above. Between two genes *i*_1_, *i*_2_ ∈ **X**, 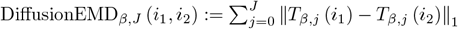, where 0 *< β <* 1*/*2 is a meta-parameter used to balance long- and short-range distances and *J* is the maximum scale considered here. 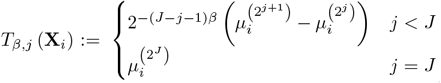, where 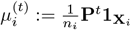 is a kernel density estimate over 𝒢_*cell*_. Graph Fourier Mean Maximum Discrepancy (GFMMD) [62] is defined via an optimal witness function that is smooth on the graph and maximizes the difference in expectation between the pair of gene distributions.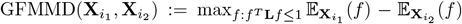, holding for any construction of a positive semidefinite Laplacian matrix **L** and chosen threshold *T* = 1.

For these approaches, multiscale signal features 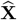 are computed prior to distance calculation. We reduce the dimensionality of these features via an autoencoder *D*○*E*, where 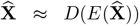 and 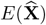 is the embedding.

#### Computing eigenscores

Eigenscores were proposed as a topologically motivated mathematical method for feature selection, and they were also shown to be useful for mapping the gene space to distinguish cell types [44]. Eigenscores rank signals or genes based on their alignment to low-frequency patterns in the data, identified through spectral decomposition of the graph Laplacian. Specifically, given the first *r* left eigenvectors of the normalized Laplacian (where *r* ≪ *n* to preserve low-frequency patterning), 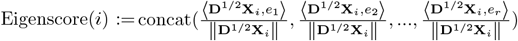. We let Eigenscore(**X**), of shape *m* x *r* represent the eigenscores for each gene *i* in **X**. We finally reduce the dimensionality for gene space mapping via an autoencoder *D* ○ *E*, where Eigenscore(**X**) ≈ *D*(*E*(Eigenscore(**X**))) and *E*(Eigenscore(**X**)) is the embedding.

#### Co-embedding of cells and genes

Finally, recent approaches incorporate cell-cell affinities through simultaneously learning embeddings for cells and genes. This methodology has the benefit of *learning* the pairwise similarities between cells, rather than constructing the cell-cell graph *a priori*, and training this module in tandem with gene-gene similarity training. siVAE [19] is a neural network consisting of cell-wise and gene-wise encoder-decoders. The cell-wise encoder takes each cell’s measurement across all features and maps cell embeddings similarly to a classical VAE, which computes an approximate posterior distribution over the location of the cell. The gene-wise encoder takes a gene’s measurement across all cells and maps gene embeddings. The decoders of both VAEs combine to output the expresion level of each feature in each particular cell, ensuring that each mapping has semantic structure. SIMBA [15] constructs a heterogeneous graph, where the nodes are cells and genes, and edge type are determined through expression level. SIMBA first bins the continuous gene expression values into a discrete distribution that preserves the shape of the original distribution, then encodes different bins as different relation types. A node embedding for each node in the graph is then learnt via stochastic gradient descent optimization of a link prediction objective. For both procedures, we evaluated only the gene space embeddings in our comparisons.

### 6.3 Diffusion wavelets and comparison to MAGIC

Diffusion wavelets, as compared to diffusion maps [21] and related approaches, such as MAGIC, perform multiscale analysis of graphs and functions on graphs. By representing the cell-cell graph at multiple scales, diffusion wavelets are able to decompose low-frequency and medium-frequency components of a signal, in addition to removing noise, whereas MAGIC acts as a low-pass filter and only maintains the low-frequency components (Extended Data Figure 3).

Additionally, diffusion wavelets represent the local and global geometry of the cell-cell graph, which enables the representation of the distance between signals that are far apart. We demonstrate this with an experiment using a linear simulated trajectory with noise (also used in Figure 2 and Figure 3) (Extended Data Figure 4a). This trajectory defines our cell-cell graph, and we simulate signals on this graph as diracs defined on each cell in the trajectory. Each signal naturally has a label associated with it - the pseudotime value of the cell it was defined on (Extended Data Figure 4b). Notably, as each signal is not overlapping, approaches that do not use the cell-cell graph would not be able to capture meaningful distances between signals.

To test the signal embeddings, we regress the pseudotime label from the latent space (Extended Data Figure 4c). We hypothesized that methods that locally smooth the signal would perform well when signals are close together, as local geometry sufficiently captures pseudotime, and poorly when the signals are far apart. Therefore, we evaluated the prediction with Spearman correlation and increased separation between signals. As expected, MAGIC worsens in the performance as signals are increasingly spread apart, whereas GSPA and GSPA+QR do not show a trend related to the separation between signals (Extended Data Figure 4d). We also perform unsupervised evaluation of the embeddings at spacing 2^6^ via the correlation between the Fiedler vector of the signal graph and the pseudotime label. GSPA and GSPA+QR show high correlation versus MAGIC (Extended Data Figure 4e), and also show a qualitative association with the pseudotime label when visualized versus MAGIC (Extended Data Figure 4f).

### 6.9 Training Details

#### Default GSPA hyperparameter selection and training details

The cell-cell graph was built with PHATE using default parameters (*k*=5 and *α* decay=40) from the PCA space, as common for cell-cell graph construction. The power was set by default to 2 to mimic the dyadic scales in [22] and J was set by default to *log*(*n*) based on [93] (see Lemma 1 and surrounding discussion above). For GSPA+QR, the epsilon parameter was set to 1*e*-3. The data was first dimensionality reduced with PCA to 2048 components (which captures the majority of variation), and then an autoencoder nonlinearly reduced the dimensionality further to latent dimension of 128. The autoencoder was designed with 2 layers with bias in the encoder and decoder, with a relu activation function between layers. The models were trained for an MSE objective with an Adam optimizer with learning rate of 0.001 for 100 epochs, with early stopping (patience of 10) using the loss of a validation set 5% of the size of the training set. For all analyses, signals are first L2 normalized before projection.

#### Comparison hyperparameter and training details

For method comparisons in Figure 2 and Figure 3, we ran each method three times, including reconstructing the graph with new seeds. All signals were first L2 normalized, and, where applicable, dimensionality reduced using PCA with 2048 components and an autoencoder (AE) with latent dimension of 128 (PCA+AE; same configuration as for GSPA). For raw measurements, we ran PCA+AE on **X**. For MAGIC(**X**), we compute the diffusion operator with default parameters. We then project the signals onto this diffusion operator and run PCA+AE. We compute eigen-scores based on the approach described in [44], then dimensionality-reduced with PCA+AE. We learned multiscale representations with DiffusionEMD and GFMMD, then dimensionality reduced with PCA+AE. For signal-signal graphs, *k*-NN graphs were generated from the signals with *k* = 5. Node2Vec was run on this graph with latent dimensionality of 128, walk length of 80, and 10 walks. GAE_no-att_ was run with graph convolutional layers, and GAE_att_ was run with graph attention layers on this graph. The GAE configuration matched the previous AE configuration. For SIMBA, we constructed a heterogeneous cell-gene graph using default parameters, without highly variable genes. We then trained the graph embedding with 128 dimensions, auto-estimating weight decay. For siVAE, we constructed the encoder-decoder architecture with the same number and size of layers as our GSPA autoencoder. We additionally employed 2000 iterations, *mb*_*size*_ of 0.2, *l*2_*scale*_ of 1e-3, learning rate of 1e-4, decay rate of 0.9, and early stopping with a patience of 100 iterations. We used relu activations in between layers.

### 6.10 Datasets and Pre-processing

#### Simulated datasets with Splatter

Three datasets were simulated using Splatter [106] with one (linear) trajectory, two branches, and three branches. All datasets were simulated with 10,000 cells and 10,000 genes, where cells were distributed equally between branches (where applicable). The dropout probability was set to 0.95 to generate “noisy” datasets, and each dataset had associated “true” noiseless counts from the same experiment. After simulation, genes expressed in less than 50 cells were removed, and the matrix was L1 normalized for library size and square-root transformed (or log-transformed for robustness analysis in Extended Data Figure 10). This resulted in 8821 genes in the linear simulation, 8820 genes in the two-branch simulation, and 8823 genes in the three-branch simulation. Cells were then visualized with PHATE.

#### Peripheral blood mononuclear cell (PBMC) dataset

This data consisted of 2,638 cells and 1,838 genes, following the `scanpy` preprocessing workflow (https://scanpy-tutorials.readthedocs.io/en/latest/pbmc3k.html) to analyze 10x Genomics data acquired from 10x Genomics [1]. Cells with fewer than 200 genes expressed and genes expressed in fewer than 3 cells were removed. Cells with over 2500 total counts or over 5% mitochondrial counts were removed. The data was L1 library size normalized and log-transformed, and highly variable genes were preserved and scaled.

#### Embryoid body (EB) dataset

This data was derived from [69] and captures cellular populations within the embryoid body differentiation process. We followed the preprocessing procedure from the original work, removing cells with library size higher than the 75% and lower than the 20% for each sample. Genes expressed in fewer than 10 cells were removed, and the data was L1 library size normalized. The top 10% of cells with highest mitochondrial expression were removed, and the data was square-root transformed. This resulted in 16,821 cells and 17,845 genes.

#### Three-timepoint scRNA-seq dataset (previously unpublished)

Mice were infected with lymphocytic choriomeningitis virus (LCMV) Armstrong (Acute) and Clone 13 (Chronic), and CD8+ CD44+ Tetramer+ T cells were FACS sorted prior to 10X Chromium 5p single-cell RNA sequencing at day 4, day 8, and day 40. 3-5 mice were infected for each timepoint/condition in a staggered manner to enable same day take down of each timepoint. Spleens from mice were pooled for each timepoint/condition and sorted prior to their loading on the Chromium instrument. 10,000 cells were loaded into a lane of the instrument for each timepoint/condition. The resulting 10X libraries were sequenced on an Illumina NovaSeq with an approximate read depth of 20,000 reads per cell. We then processed the data using Cell Ranger [108] before further filtering. Cells expressing less than 200 genes, with less than 500 counts or more than 25000 counts, were removed. Genes expressed in less than 3 cells were removed. Cells with mitochondrial percentage greater than 6% were removed. We then L1 normalized for library size, log-transformed, and clustered cells using Leiden clustering [94], removing contaminating populations enriched for non-CD8+ T cell markers. The acute and chronic datasets were combined, and highly variable genes were detected as the top 10% of genes using `scprep` (https://scprep.readthedocs.io/en/stable/). This resulted in 14,152 genes and 39,704 cells detected across datasets, with 6,811 cells from Acute Day 4; 7,418 cells from Acute Day 8; 6,740 cells from Acute Day 40; 6,205 cells from Chronic Day 4; 7,553 cells from Chronic Day 8; and 4,977 cells from Chronic Day 40. The combined datasets were then visualized with PHATE, and key marker genes were visualized on the PHATE embedding with MAGIC. Graphs for PHATE and MAGIC were built with default parameters, except *k* for the *k*-NN graph construction was set to 30 due to the larger number of cells.

#### Transcription factor perturbation scRNA-seq dataset (previously unpublished)

The sgRNA library was cloned into an MSCV retroviral backbone containing an expression cassette for human CD2 as a selectable marker. The library targets 39 genes the majority of which are transcription or epigenetic factors with three unique sgRNAs per target gene. sgRNA sequences were chosen via CHOPCHOP. Negative control sgRNAs were spiked in to make up 15 percent of the final library. Retrovirus production was performed using platinum-E cells. P14 Cas9 transgenic CD8+ T cells were isolated (StemCell Mouse CD8 Selection Kit) and activated with anti-CD3/CD28 activation beads (Dynabeads Mouse T activator) for 1 day prior to infection with sgRNA library retrovirus using retronectin and protamine sulfate. The cells were then grown in human IL-2 (5ng/mL) for 2 days prior to magnetic selection to enrich for transduced cells based on the human CD2 selection marker (StemCell Human CD2 positive selection kit). 100,000 cells were then transferred into 7 day 1 LCMV-Armstrong infected mice. At day 8 of infection, mice were sacrificed and P14 T cells were sorted from the spleens. Equal numbers of P14s were pooled from 7 mice prior to sorting and then loaded into 6 wells of a ChromiumX instrument to perform paired 5’ scRNA and CRISPR sequencing. 40,000 cells were loaded into each well to recover approximately 20,000 cells per well. The resulting 10X libraries were sequenced on an Illumina NovaSeq at a read depth of 20,000 reads per cell for RNA and 5,000 reads per cell for CRISPR feature barcode. The data was then processed using Cell Ranger [108] prior to downstream analyses. We removed cells with less than 200 genes detected and cells with less than 500 counts and more than 20000 counts. Cells with mitochondrial percentage greater than 5% were removed. To remove doublets, the data was log-transformed, scaled, and centered, and doublets were identified with DoubletFinder [67] with parameterSweep v3 and expected doublet percentage of 7.5%. We then clustered cells using SNN-based Louvain clustering [85] and removed contaminating populations enriched for non-CD8+ T cell markers. We further filtered the cells to annotate their pertubation identity by assigning an sgRNA identity to a cell if there was *>*1 UMI for that sgRNA and it made up *>*80% of the sgRNA reads in that cell. Finally, we integrated the replicates with CCA [53] using the top 2000 highly variable genes and removing TCR-related, mitochondrial, proliferation, and immunoglobulin genes. This resulted in 23,206 cells and 1,795 genes across all perturbations.

#### Peripheral tolerance scRNA-seq dataset (Damo et al.)

We obtained scRNA-seq data from [24] and pre-processed it as in the scRNAseq Methods section. There were 21,178 cells and 21,515 genes after preprocessing. This corresponds to 8,167 cells from Ag ON samples; 3,944 cells from Ag ON/CPI samples; and 9,067 cells from Ag OFF (i.e. no AG) samples. Cells were visualized with PHATE and key genes were visualized on the PHATE embedding with MAGIC. The cell-cell graph for PHATE, MAGIC, and GSPA were built with default parameters, except *k* for the *k*-NN graph construction was set to 40, and *α* was set to 10 to match the analysis in the original work.

#### 10x Human Lymph Node Spatial Transcriptomics Dataset

10x Genomics data was obtained from the 10x website [2] and downloaded via the `scanpy` package [101]. According to their website, 10x obtained fresh frozen human lymph node tissue from BioIVT Asterand Human Tissue Specimens. The tissue was embedded and cryosectioned as described in the Visium Spatial Protocols Tissue Preparation Guide (Demonstrated Protocol CG000240). Tissue sections of 10 µm thickness were placed on Visium Gene Expression Slides. We removed spots with less than 5000 counts, more than 35000 counts, and over 20% mitochondrial counts. We removed genes detected in fewer than 10 cells, L1 normalized for library size, and log-transformed the data. After the above pre-processing, there were 3861 spots and 19,685 genes detected. The top 2000 highly variable genes were selected following the `scanpy` tutorial for this dataset.

#### Immunotherapy response in melanoma patients scRNA-seq dataset (Sade-Feldman et al.)

We obtained pre-processed scRNA-seq data with annotated cell types and other relevant metadata (e.g. sample labels, patient response) from [77] and the Single Cell Portal (https://singlecell.broadinstitute.org/single_cell). From this data, there were 48 samples, which corresponds to 19 pre-therapy samples and 29 post-therapy samples, as well as 31 nonresponder samples and 17 responder samples. There were 15,300 cells and 12,364 genes detected across all samples, with 10,190 cells from nonresponders and 5,110 from responders.

### 6.11 Computational Details

#### Diffusion wavelets versus MAGIC

For Extended Data Figure 3, the smooth signal was defined based on visualization coordinates to capture the low-dimensional axis of variation. The oscillating signal was defined by the sine of 4 times the smooth signal to define a medium-frequency component. The noise was generated based on random samples from a uniform distribution between [ − 1, 1). The aggregate signal was defined based on the sum of these three components. The denoised signal was computed with MAGIC and the wavelet dictionary was computed with GSPA and *J* = 5.

#### Dirac signals comparing GSPA and MAGIC

For the dirac experiment (Extended Data Figure 4), we simulated and pre-processed a linear trajectory of cells as described above, and we defined signals as a dirac on each cell, where the signals naturally had a pseudotime label based on the cell it was defined on. We learned unsupervised embeddings for GSPA+QR, GSPA, and MAGIC as described above. Then, we increased the distance between signals by subsampling every other cell in the pseudotime-ordered trajectory, then every 2^2^ cells, every 2^4^ cells, and so on, and we learned embeddings for the signals increasingly spread apart. We then evaluated the embeddings by repeated K fold splits and ridge regression to predict the pseudotime labels, comparing methods based on Spearman correlation between prediction and true labels across 10 runs. Gene embeddings were visualized with PHATE.

#### Simulated coexpression experimental details

For the coexpression experiment with a linear trajectory (Figure 2) and two and three branches (Extended Data Figure 5), we generated simulated data as described above, then defined signals as the gene features from the simulation experiment. Because of the simulation design, this meant we have both noisy **X** and noiseless **X**^*′*^ versions of the same gene signals. This allows us to compute “ground truth” coexpression as the Spearman correlation between all noiseless pairs of genes. Given the large number of genes and the nature of biological data, the large majority of gene-gene pairs had a near-zero correlation. The correlation also was associated with the library size of the genes in the pair. Therefore, we stratified the labels based on correlation and the mean library size of the pair within each correlation bin. We learned unsupervised gene embeddings for all comparisons as described above, then, for an equal number of pairs per stratification bin, we computed the distance between gene embedding pairs and the (anti-)correlation with the true coexpression.

To identify gene modules, we visualized the GSPA+QR embedding with PHATE and used Leiden clustering. To map these gene modules back to the cells most enriched for the modules, we leveraged a gene set enrichment approach from [78] and implemented in [101]. This approach provides a score defined on all cells as the average expression of a set of genes subtracted with the average expression of a reference set of genes. Using the 25% of the number of genes as the number of bins, we calculated a cell enrichment score for each gene module and visualized this enrichment score versus pseudotime.

To analyze how the gene embeddings relate to peak over time (trajectory analysis), we binned cells into 100 bins based on pseudotime values, and computed the mean expression of each gene over the binned timepoints. Then, we annotated each gene based on which bin it “peaked”, or had a maximum value. We colored all gene embeddings based on this score.

Finally, to perform archetypal analysis of the gene space, we ran Archetypal Analysis Network (AAnet) [26, 100] on the gene embeddings outputted by each method with *n*_*at*_ = 2. Then, we identified the 50 genes nearest to each archetype and used the same gene set enrichment approach as above [78,101], now visualizing the cell enrichment score on the embedding and over pseudotime for only archetypal genes, rather than all gene modules.

### Simulated localization experimental details

#### Generating simulated signals with known localization scores

For the localization experiment with a linear trajectory (Figure 3) and two and three branches (Extended Data Figure 9), we generated simulated data as described above. However, instead of using the genes as signals, we designed signals with “ground truth” localization labels (Extended Data Figure 8). We intuited that more localized signals are not defined by where they are enriched in the trajectory, but rather by how spread out that enrichment is. Thus, we aimed to constrain the size of the region where each signal could be defined, termed “window”, where the window can be defined anywhere on the trajectory and is only defined by its size.

To generate signals and associated localization scores with these properties, we used the ground truth pseudotime label (provided by Splatter) scaled to be between 0 and 1, and we defined window size *δ*. Then, we randomly selected a timepoint *t* between [*δ/*2, − 1 *δ/*2] and defined a pseudotime window [*t*− *δ/*2, *t* + *δ/*2]. Next, we sampled 500 cells from all cells within this pseudotime window, and we let the signal equal 1 on these cells and 0 on all other cells (Extended Data Figure 8).

For each of five window sizes *δ*∈ { 0.2, 0.4, 0.6, 0.8, 1.0}, we generated 50 signals, resulting in 250 signals total. As smaller *δ* corresponds to a higher localization score, we defined the true localization score for each signal to be 1 − *δ*. This score is unrelated to where the signal is defined based on randomly selected *t*. Furthermore, all signals are defined on exactly 500 cells, so the localization score is not associated with the number of cells expressing the gene.

#### Predicting localization from uniform signal

We then defined a uniformly-enriched gene as a signal equal to 1 on all cells. L2 normalizing all signals, we embedded the uniform gene with other signals and computed the (multiscale) distance of all signals to the uniform distribution. We intuited that the uniformly expressed gene should be closest to signals with a low localization score (*δ* = 1.0) and furthest from signals with a high localization score (*δ* = 0.2). That is, the distance to the uniform signal can be considered the predicted localization score (as described in Definition 2). We therefore predicted the localization score for all signals and evaluated methods based on Spearman correlation of predicted and true localization.

#### Computing localization for comparisons

For GSPA+QR, GSPA, and MAGIC, this involved projecting the uniform signal onto the cell representation/dictionary and calculating the distance between the projected uniform signal and all other projected signals. Eigenscore and GFMMD defined a version of this localization based on the L2 norm of their embeddings, so we evaluated localization using this measure. For DiffusionEMD, we learned a multiscale representation of the uniform signal, and we computed the distance to all other signals before dimensionality reduction. For the raw measurements, we took the distance of the uniform signal to all other signals before dimensionality reduction. For Node2Vec and the GAE approaches, we built a signal-signal graph with the uniform signal and embedded these graphs, then computed the L2 distance between the uniform embedding and the other signals. For SIMBA and siVAE, which learn a low-dimensional representation of the genes directly, we learned a low-dimensional embedding of the uniform signal and computed the distance to all other signals in this latent space.

#### Robustness to transformation and graph construction details

For robustness analysis, we followed the same training details as in the previous section, only changing the input data transformation (`log` and `sqrt`), run seed (two runs), choice of *k* for *k* nearest neighbors graph construction (5, 15, 25, 50), and choice of graph construction for nearest neighbors graph (*k* nearest neighbors (*k*NN, shared nearest neighbors (SNN), and graph construction with an adaptive *α*-decaying kernel, as previously described).

#### Batch effect experimental details

To analyze the effects of batch effect and batch effect correction with GSPA, we simulated a dataset of 2000 cells and 10,0000 genes using Splatter [106], with 1000 cells in each batch, 3 clusters with equal probability, 0.6 DE probability (probability that a gene will be selected to be differentially expressed), and batch factor location and scale of (0.1, 0.01) (where batches are specified by generating a small scaling factor for each gene in each batch from a log-normal distribution). This data was then processed to remove genes expressed in fewer than 50 cells, L1 normalized for library size, and log transformed. We then computed gene embeddings with default GSPA+QR for all genes, coloring by DE factor and batch effect factor. To correct the graph for batch effect, we constructed a mutual nearest neighbors (MNN) graph, introduced in [42], and reran and revisualized PHATE and GSPA+QR with the corrected graph. We then computed cell type association scores for each of the annotated clusters and visualized those scores.

#### Characterization of PBMC gene embedding

For coexpression analysis (Figure 2g), we embedded all highly variable genes using the default GSPA+QR approach. Then, to visualize cell type-specific genes, we retrieved all annotated cell type markers from PanglaoDB [31] for T cells, Monocytes, B cells, NK cells, Dendritic cells, and Megakaryocytes. We then subsetted this list to “canonical markers” (defined by PanglaoDB) that were highly variable and had highest mean expression in the correct annotated cell type in our dataset.

For analysis of localized genes (Figure 3d), we computed the localization score for all mapped genes and identified the genes with the top 25% localization score as “predicted localized genes”, and genes with the bottom 25% localization score as “predicted non-localized genes”. We then constructed the cell-cell graph using only these genes and reran PHATE. Next, we calculated the pairwise geodesic distances between all cells in the full graph and the feature-selected graph. As a proxy for the preservation of the cell-cell relationships with selected features versus all genes, we randomly chose 100,000 pairwise distances (in two runs) and computed Spearman correlation between the full graph distances and the feature-selected graph distances.

#### Characterization of EB gene embedding

For coexpression analysis (Figure 2g), we embedded all measured genes except mitochondrial genes, as they displayed a very different trend than the other genes and embedded distinctly. We used the default GSPA+QR approach. Then, to perform gene trajectory analysis, we computed the diffusion map for these genes. Based on the lineage analysis done in [69], we identified diffusion map component 4 as associated with the hemangioblast lineage, and colored the gene embedding by this component, annotating various key genes along this gene trajectory.

For analysis of localized genes (Figure 3d), we repeat the procedure as for the PBMC data to compare the full graph and feature-selected graph distances and embedding.

#### Computing cluster rank for localization versus cluster rank comparison

For the comparison in Figure 4d, we performed Leiden clustering on the cells, which identified 9 cell clusters. Using `scanpy`, we ranked genes based on the Wilcoxon rank sum test to identify genes differentially enriched in each cluster. This results in a z-score underlying the computation of a p-value for each gene for each group. For each gene, we stored the maximum score across clusters, and we ranked genes based on this maximum score. This ranking reflects how enriched the gene is, which we compare against the computed localization score.

#### Computing gene module enrichment per condition

Genes were visualized with PHATE and modules were identified via Leiden clustering (Figure 4b). Using the same gene module enrichment approach as in the simulated coexpression experiment, now using the highly variable genes as the reference set, we calculated a cell enrichment score for each gene module. As positive values of this score indicate an enrichment over the reference set, we counted cells with a score *>* 0 from each sample and normalized this count by the number of cells from each sample. This represents the gene module enrichment of cells per condition (Figure 4e).

#### Computing and comparing type 1 interferon signaling signature

To determine the enrichment of the type 1 interferon signaling gene signature, we constructed gene embeddings for all approaches designed to map the gene space. Then, we identified gene modules using Leiden clustering, chose the gene module containing canonical type 1 interferon marker *Irf7*, and selected the top 10% localized genes within the gene module. This allowed us to choose genes that were both related to type 1 interferon signaling, through similarity to *Irf7*, but were also unbiasedly selected based on the calculated gene modules and localization score. We next wanted to add additional comparisons to other canonical approaches for identifying gene signatures. To compare against analysis done by clustering cells and identifying differentially expressed genes, we selected the top 100 DEGs from each cell cluster (obtained as noted above). Finally, to compare against factor analysis approach cNMF, we extracted the gene program for which *Irf7* had the highest loading, then selected the genes with the highest 10% loading score to that program. To compare the biological relevance of selected genes from each comparison, we performed gene set enrichment analysis using Enrichr [103] and the BioPlanet gene set resource [45], and visualized enrichment scores for two type 1 interferon-related gene sets.

#### Building module-specific gene coexpression networks

To build module-specific gene coexpression networks (Figure 4f), we identified the top 10% localized genes in each gene module, then built a *k*-NN graph with *k* = 5 from the GSPA+QR gene representations. Networks were then visualized with Cytoscape [80].

#### Building perturbation-specific gene coexpression networks

First, we identified genes that were in the top 25% localized in both the negative control and the knockout (KO). Then, we built a *k*-NN graph with *k* = 5 for the negative control genes, and a *k*-NN graph with *k* = 100 for the KO genes from the GSPA+QR representations. We subtracted the KO adjacency matrix from the negative control adjacency matrix and built a new graph from the positive entries, visualizing this graph with Cytoscape. This effectively identifies coexpression edges that are in the negative control that are not in the KO gene-gene graph. Notably, the difference in *k* was in order to emphasize connections that were very similar in the negative control and very different in the KO. For visualization, we removed disconnected subgraphs consisting of 2 or fewer nodes.

#### Calculating cell type-specific communication with CellPhoneDB

As a comparison with the canonical approach for cell-cell communication, we used the permutation test developed in [29] and implemented in `squidpy` [73]. The test was run with default parameters and the cell type annotated labels from the original work [24]. We then visualized the results for the two interactions of interest (CCL5-CCR5, PD-L1-PD-1).

#### Determining cells enriched with ligand and receptor in GSPA-LR analysis

In our analysis, we ignore cell type labels and run the GSPA-LR communication pipeline described in Figure 5a using the CellChatDB [47] intercellular communication database, which contains triplets (ligand, receptor, pathway), where the ligand is the source gene, receptor is the target gene, and pathway is an attribute defining to which communication pathway the interaction belongs. LR pairs were embedded with PHATE, and modules were identified via Leiden clustering (Figure 5e). Then, to map these gene modules back to the cells most enriched for the modules, we leveraged the gene set enrichment approach from [78] to identify cells enriched for the ligands and the receptors for each module (Extended Data Figure 14). We visualize the enrichment score for the ligands and receptors in each module, and calculate the condition-specific score as above.

#### Calculating LR module enrichment scores

For each module, we convert the ligand-receptor pairs into a list of all unique genes within the module, and compute gene set enrichment scores with this list using Enrichr [103] and the BioPlanet database [45]. Gene sets enriched for each module are ranked based on the combined score (as determined by Enrichr), and the top 5 gene sets are visualized for modules 5 and 19.

#### GSPA-multimodal for spatial transcriptomic data

To leverage GSPA-multimodal for analyzing spatial transcriptomic data, we constructed an integrated diffusion operator using the approach delineated in [58], which defines two cell-cell affinity graphs G_*exp*_ and G_*spat*_ based on expression similarity and spatial location, respectively, then constructs a diffusion operator that integrates these two graphs (see also GSPA for multiple modalities section above). Then, we build an integrated wavelet dictionary with this operator and learn gene embeddings as previously described.

#### Calculating spatial variability with SpatialDE

We compared GSPA-multimodal localization on spatial transcriptomic data with SpatialDE [87], run using default parameters. Spatially variable genes were determined as genes with *q <* 0.001 and *FSV >* 0.2 (where FSV is the fraction of variance explained by spatial variation). We visualized these spatially-variable genes on our gene embedding and colored them by FSV. We then compared the localization score of spatially variable and non-spatially variable genes with a one-sided Wilcoxon rank sums test, where *p* = 9.47*e* − 07. All genes were visualized with spARC denoised counts, but GSPA-multimodal and SpatialDE were run with the original data.

#### Building module-specific cell-state communication networks from spatial transcriptomic data

We visualize gene embeddings with PHATE and identify gene modules via Leiden clustering (Figure 6c). For each gene module, we built a *k*-NN graph (*k*=5) and kept only edges that existed in OmniPathDB, adding directionality based on the OmniPathDB annotation. We then mapped the gene signaling network to the cell-type signaling networks. We repeat the following for each gene module graph: for each directed edge (*gene*_*s*_, *gene*_*t*_), for all pairs of cell states (*celltype*_*a*_, *celltype*_*b*_), if *gene*_*s*_ is differentially expressed in *celltype*_*a*_, and *gene*_*t*_ is differentially expressed in *celltype*_*b*_, we add a directed edge from *celltype*_*a*_ to *celltype*_*b*_. We finally use Cytoscape to visualize intercellular communication edges in blue, and intracellular communication edges within the same cell type (i.e. (*gene*_*s*_, *gene*_*t*_) is intracellular and *celltype*_*a*_ = *celltype*_*b*_) in red.

#### Learning patient embeddings and immunotherapy response

For the GSPA-Pt framework, we built a combined cell-cell graph across all patients, then split the wavelet dictionary into patient-specific dictionaries and projected the gene signals for each patient onto the patient-specific dictionary. We then performed PCA with 5 components and flattened these gene representations into a single vector of size 1 × 5*m* to represent the patient. We used the first five PCs to represent the patient rather than the autoencoder embedding (as in previous analysis) because the PCs allowed for more interpretable analysis of the coefficients of the classifier. A single dimension of the latent space of the autoencoder may not necessarily capture the major axes of variation for a gene, but the first dimension of the gene PC definitionally captures the major (linear) axis of variation.

For comparison, we performed GSPA using the patient indicator signals on the cell-cell graph, then ran PCA+AE as described above. We also computed the mean expression across all genes for each patient. Finally, we computed the proportion of all clusters (representing immune cell types) and all CD8 clusters (representing CD8 cell types). Using these as unsupervised patient representations, we then classified response using a ridge classifier, comparing based on AUROC of classification. Given that the ridge classifier is a linear model, the coefficients represent the features of the patient representation most important for prediction. The features correspond to five components for each gene, so we can map the coefficients to genes relevant for prediction (Extended Data Table 5). We visualize all patient embeddings with PHATE.

## 7 Extended Data Figures

**Extended Data Figure 1.**
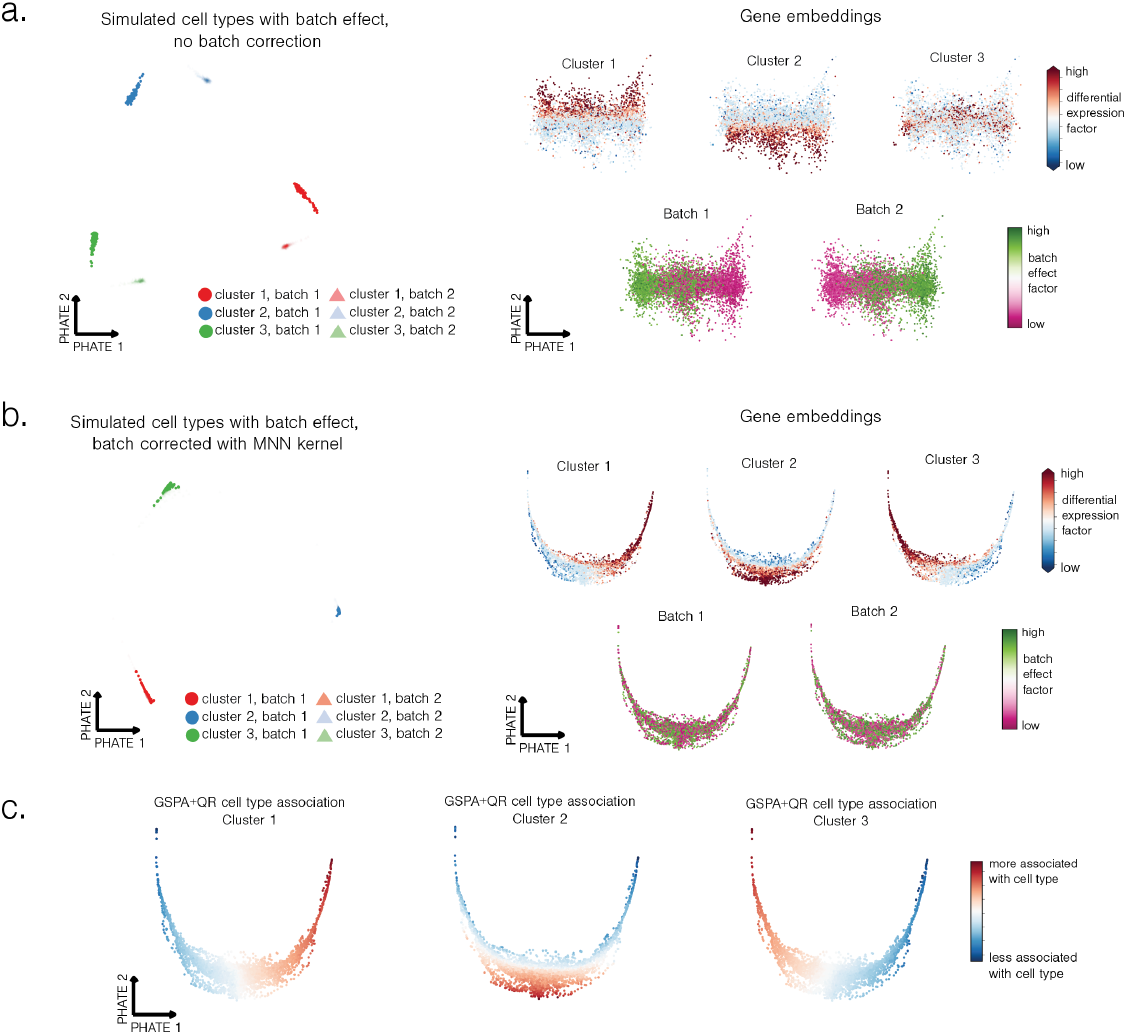
GSPA robust to batch effect. a. Dataset simulated with 3 clusters and 2 batches with batch effect. Gene embeddings colored by ground truth cluster association (differential expression factor) and batch effect association (batch effect factor) show separation by both. b. Dataset with batch effect corrected. Gene embeddings separate by cluster, but not batch effect. c. GSPA cell type association score correctly identifies relationship between genes and each cluster.

**Extended Data Figure 2.**
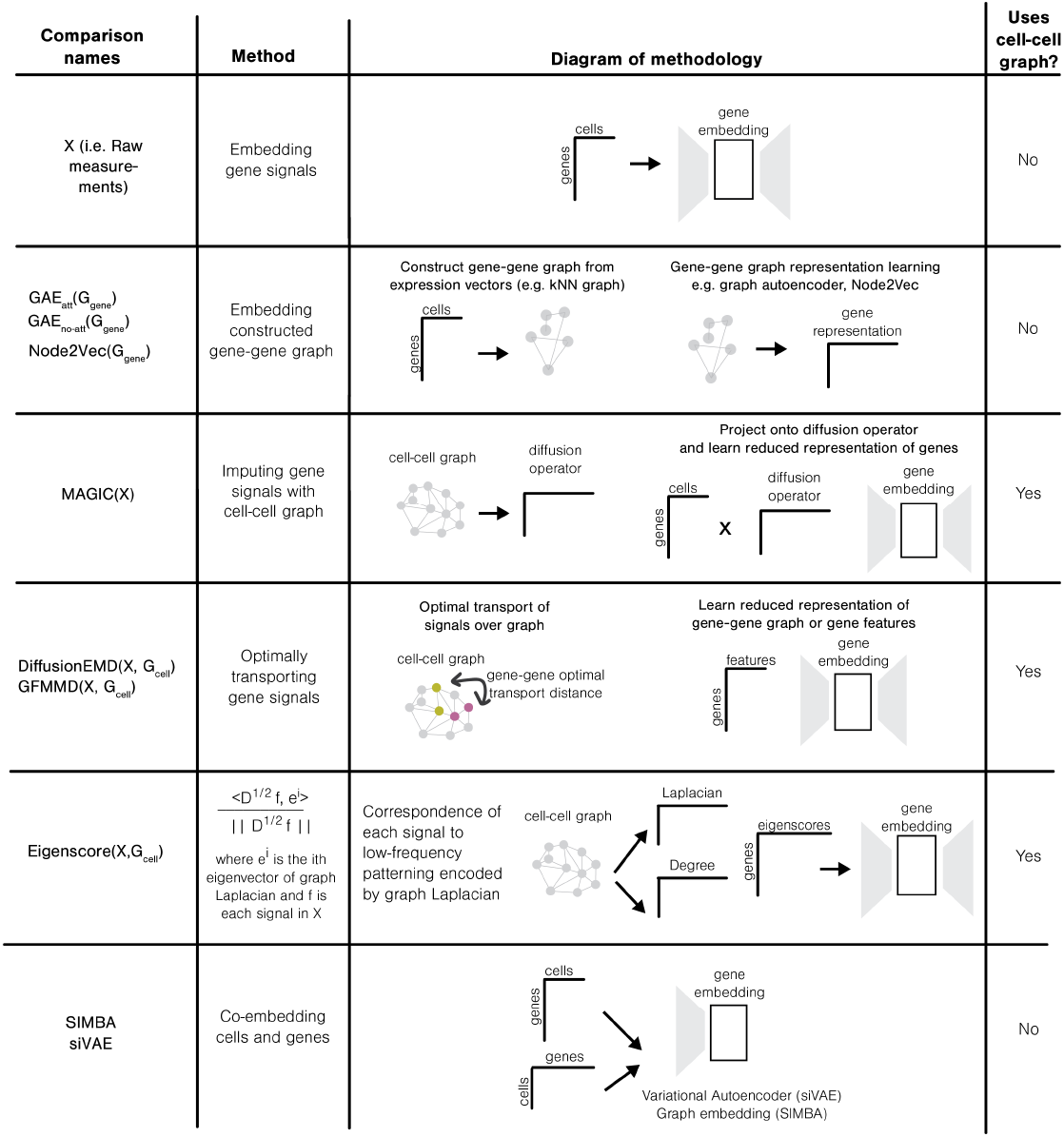
Overview of Gene Signal Pattern Analysis Comparisons.

**Extended Data Figure 3.**
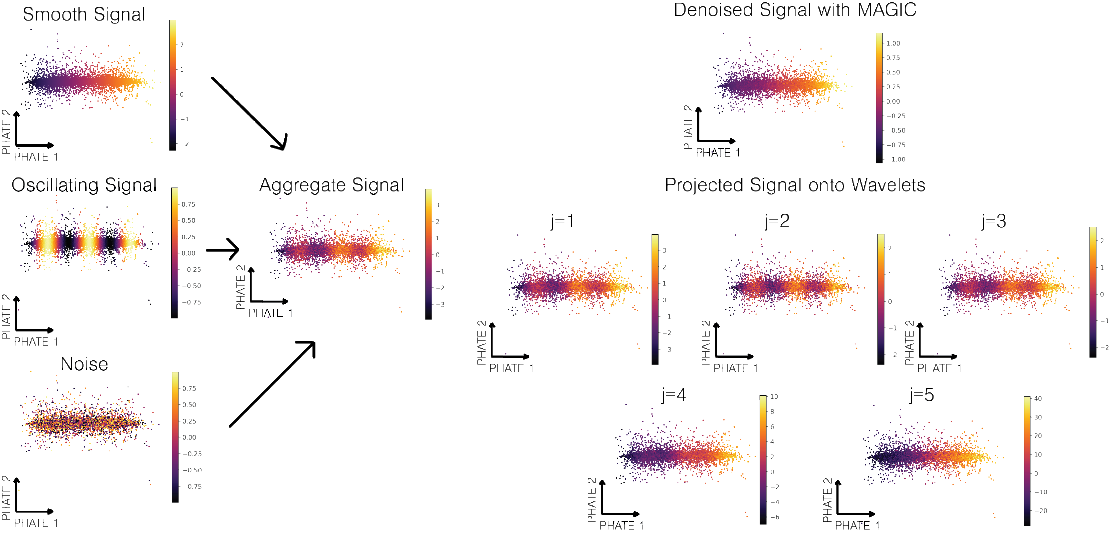
Simulated signal with low-frequency, medium-frequency, and high-frequency components. MAGIC denoises the signal and preserves the low-frequency components, whereas projection onto the wavelets reveals both medium and high-frequency components.

**Extended Data Figure 4.**
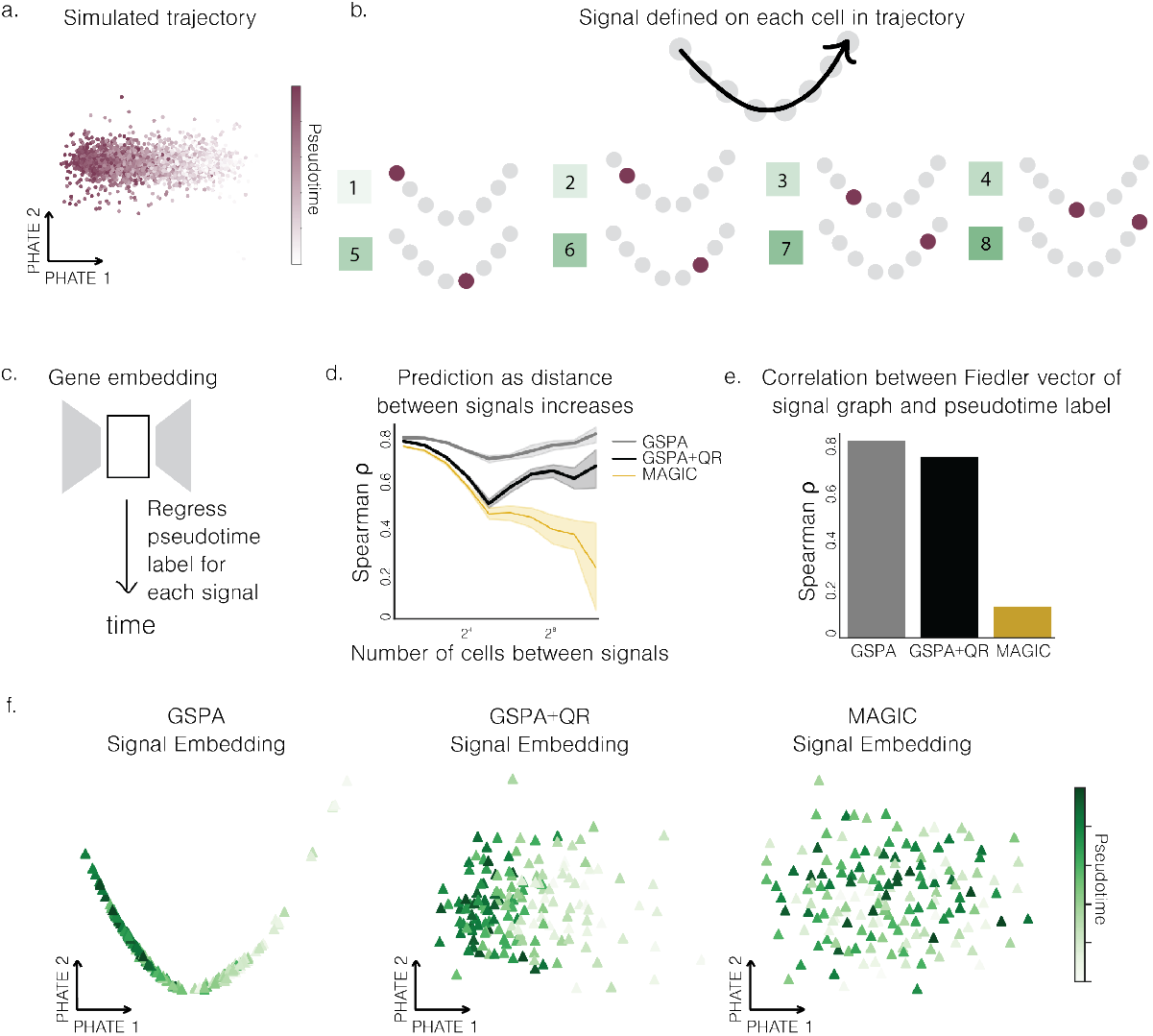
GSPA captures local and global geometry of cell-cell graph, enabling comparison of far apart gene signals. a. Simulated linear trajectory with noise visualized with PHATE. b. Schematic of generation of signals, defined on a single cell for each signal. c. Schematic of experimental setup. d. Spearman correlation evaluating performance on regression task as signals are further separated across 10 runs. e. Spearman correlation between Fiedler vector of signal graphs and pseudotime label. f. PHATE visualizations of signal embeddings colored by pseudotime labels.

**Extended Data Figure 5.**
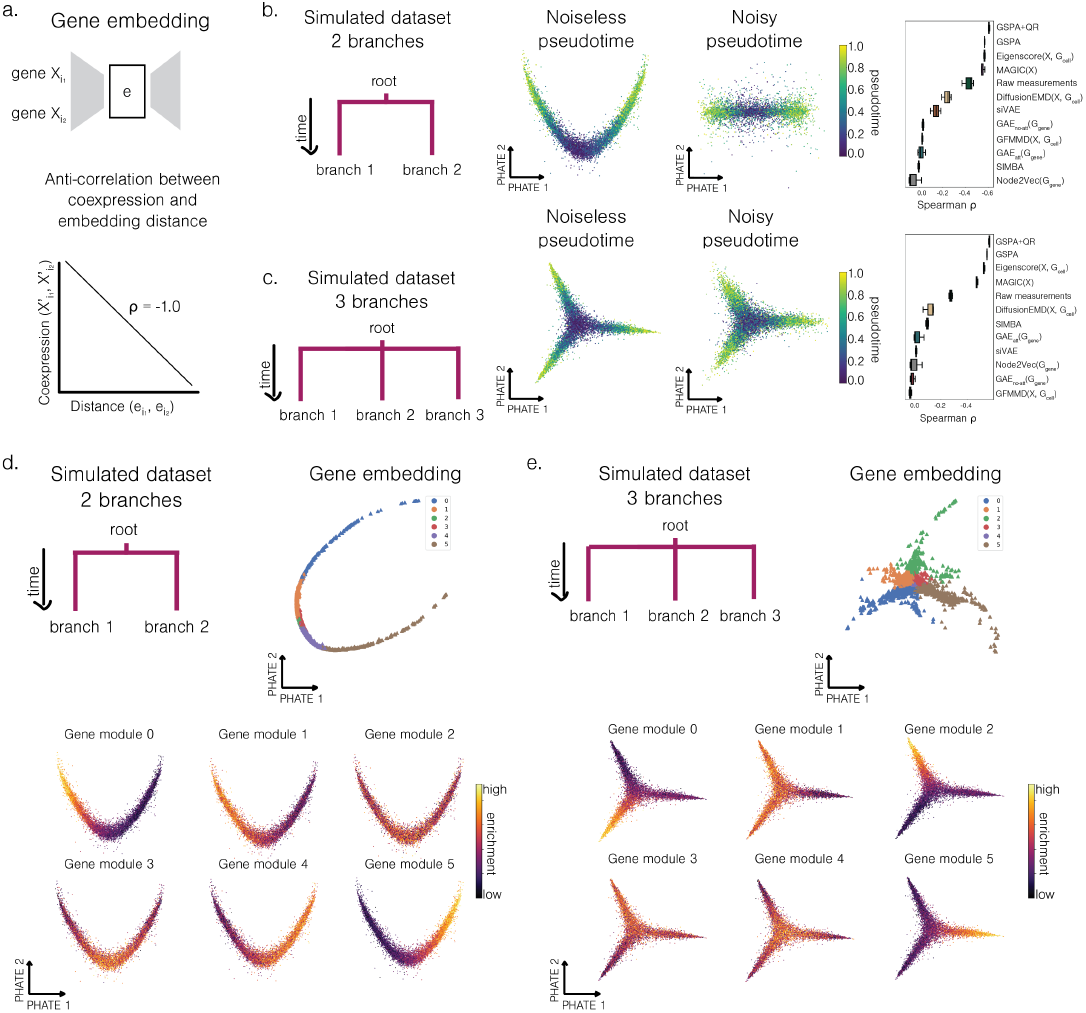
GSPA preserves coexpression in two alternative single-cell simulations. a. Experimental setup. b. Simulated dataset with two branches schematic. PHATE embedding of cells from noiseless simulation and noisy simulation, colored by pseudotime. Spearman correlation evaluating performance for all comparisons across 3 runs. c. Simulated dataset with three branches schematic. PHATE embedding of cells from noiseless simulation and noisy simulation, colored by pseudotime. Spearman correlation evaluating performance for all comparisons. d. PHATE embedding of genes from two branch simulation, colored by gene module assignments. Cells colored by gene module enrichment score. e. PHATE embedding of genes from three branch simulation, colored by gene module assignments. Cells colored by gene module enrichment score.

**Extended Data Figure 6.**
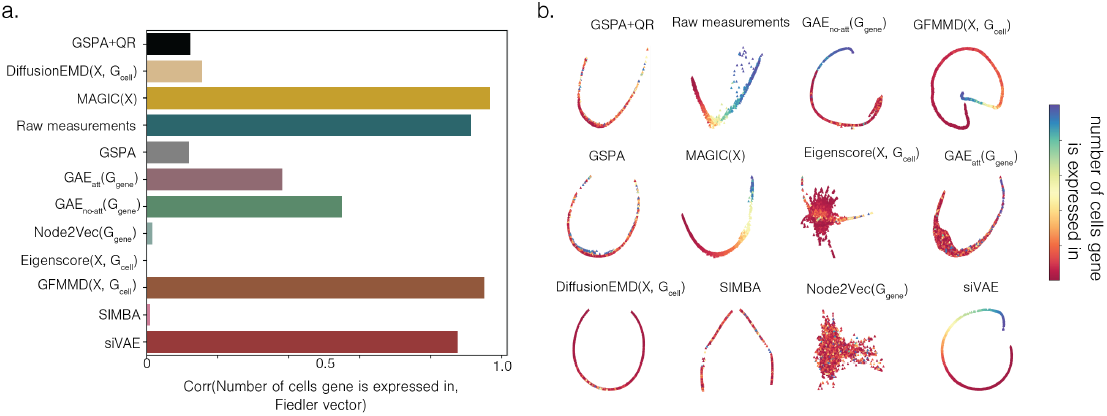
Majority of methods primarily capture number of cells expressing gene as the primary axis of variation. a. Spearman correlation between Fiedler vector of signal graphs and number of cells gene is expressed in. b. PHATE visualizations of gene embeddings colored by number of cells gene is expressed in.

**Extended Data Figure 7.**
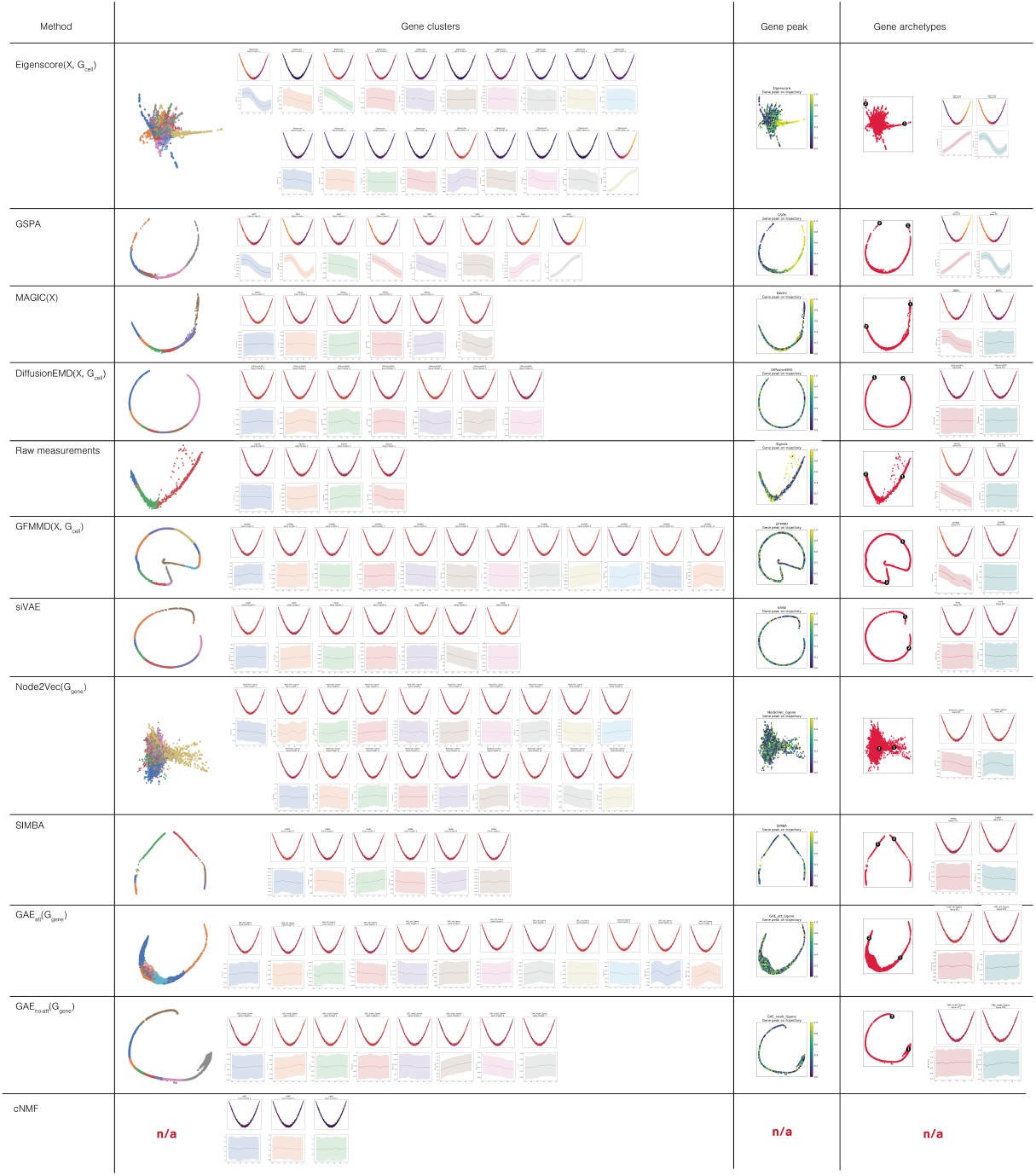
Gene embedding visualization, clustering analysis, trajectory analysis, and archetypal analysis for all comparisons plus cNMF, which enables gene module analysis only.

**Extended Data Figure 8.**
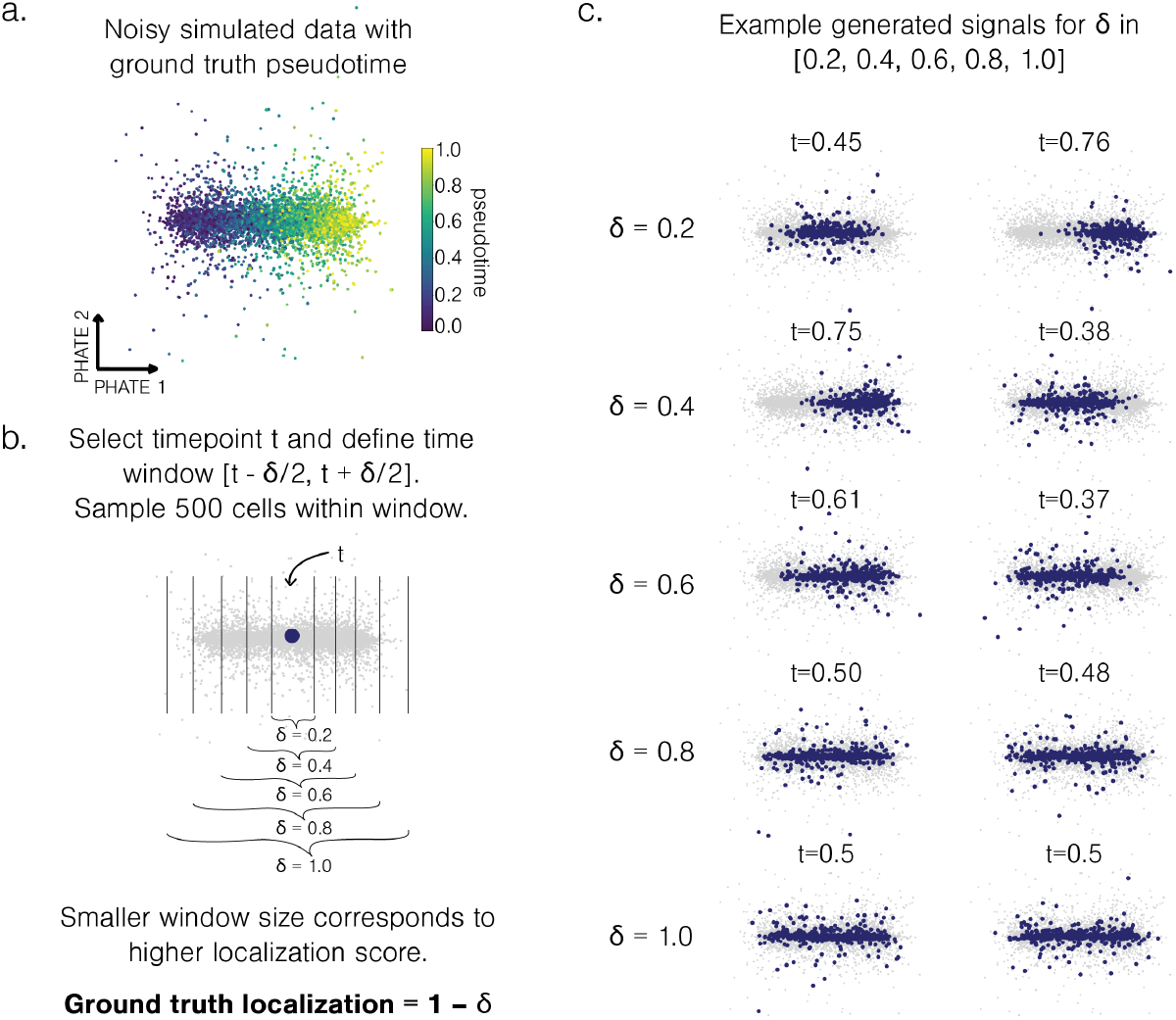
Schematic of generation of signals for localization experiment. a. Noisy simulated data with pseudotime. b. Selection of windows of size *δ*, where ground truth localization is 1 − *δ*. c. Examples of generated signals of different *δ*.

**Extended Data Figure 9.**
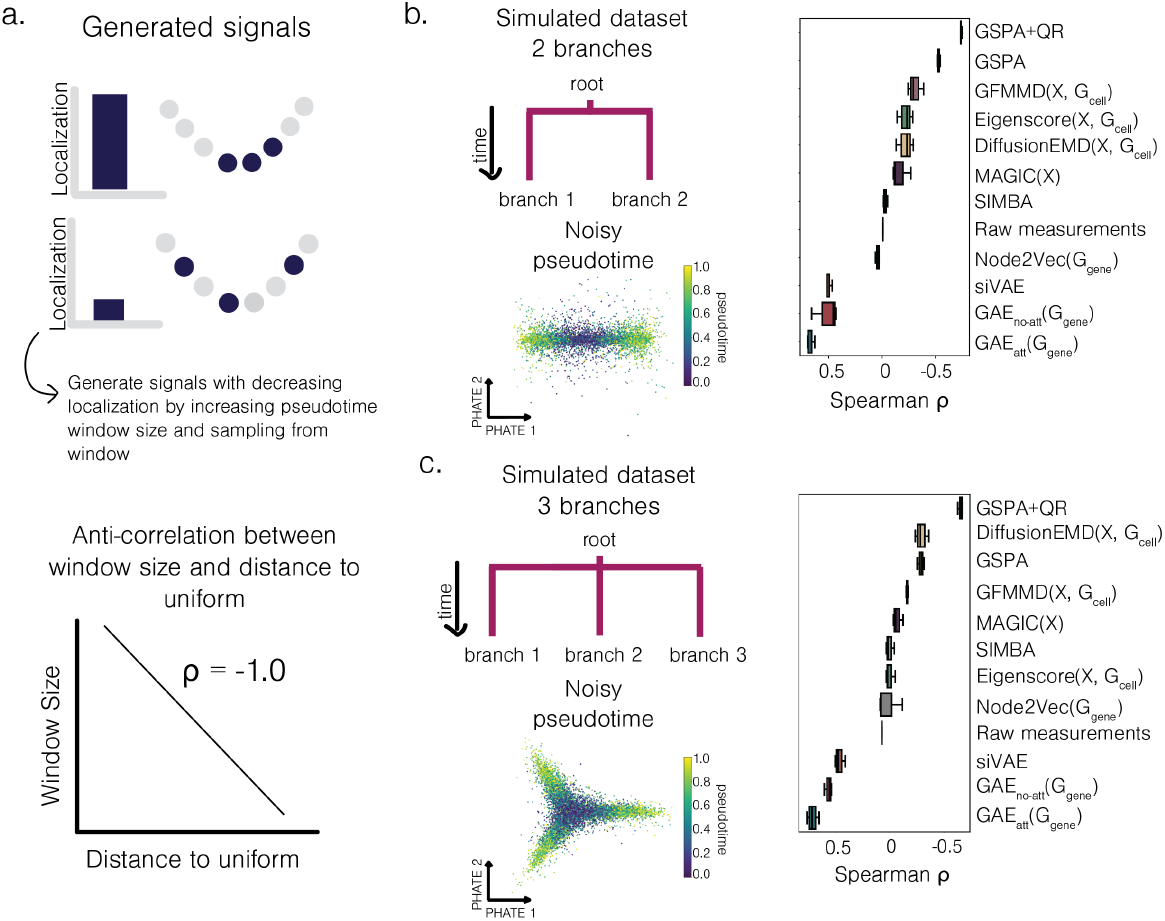
GSPA captures localized genes in alternative single-cell simulations. a. Diagram of generated signals based on pseudotime window and anti-correlation between window size and localization. Two branch noisy simulated dataset, visualized with PHATE and colored by pseudotime. Spearman correlation evaluating performance for all comparisons. c. Three branch noisy simulated dataset, visualized with PHATE and colored by pseudotime. Spearman correlation evaluating performance for all comparisons across 3 runs.

**Extended Data Figure 10.**
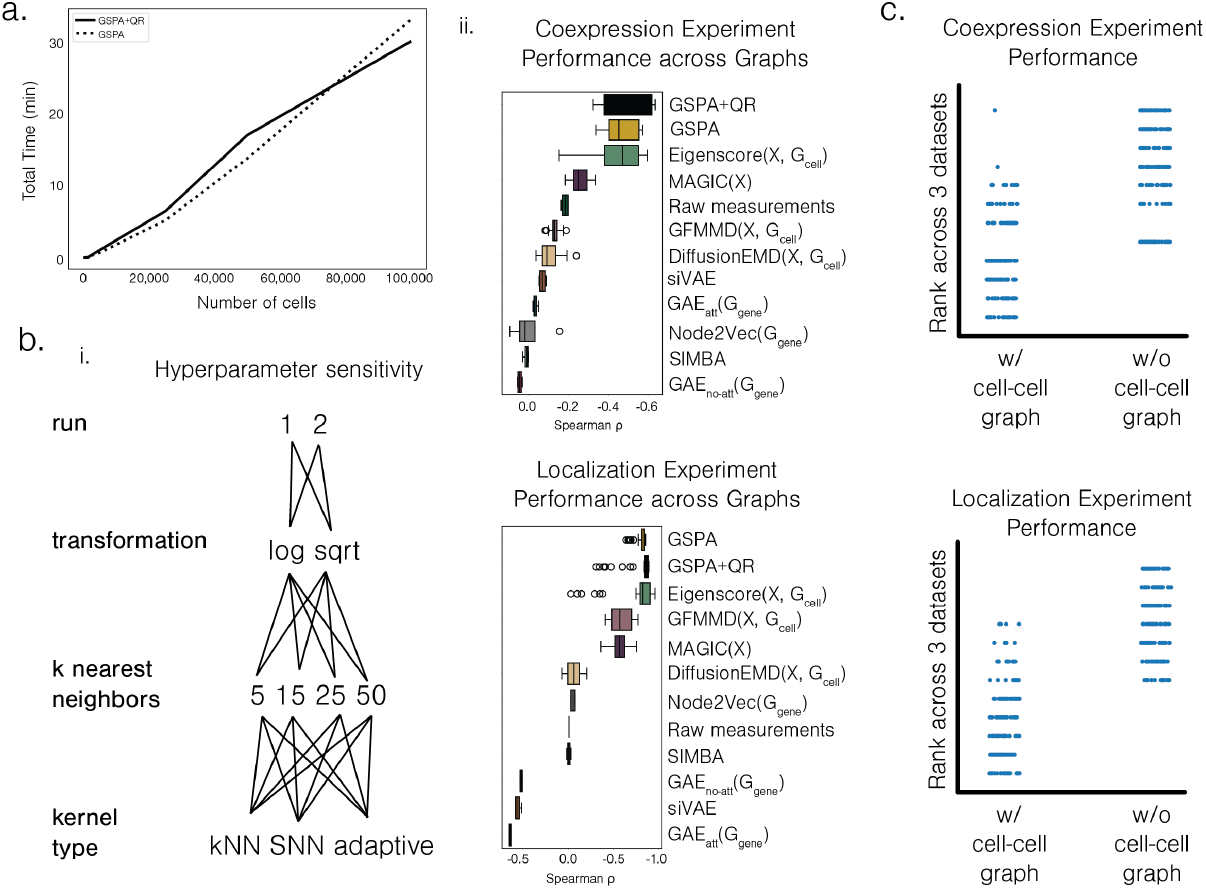
GSPA robust to transformation and graph construction. a. Runtime of GSPA and GSPA+QR. b. i. Schematic of grid search of 2 transformations, 4 kNN choices, 3 kernels, and 2 replicates (48 runs total). ii. Coexpression and localization experiment performance across all runs. c. Comparison of performance rank of methods that use cell-cell graph versus without cell-cell graph.

**Extended Data Figure 11.**
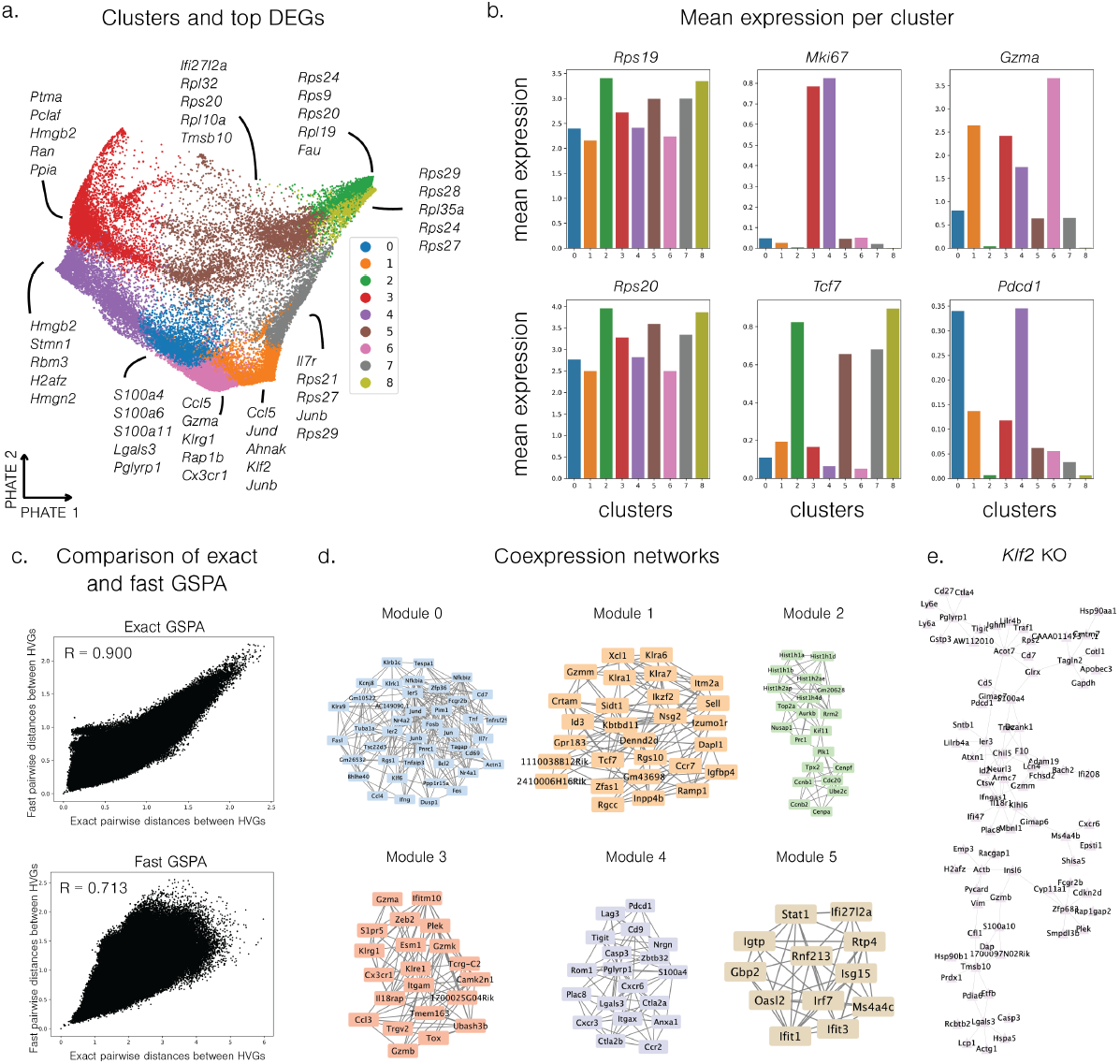
Extended CD8+ T cell analysis. a. Cells clustered with top DEGs identified. b. Mean expression of top DEGs *Rps19* and *Rps20* and key CD8+ T cell marker genes per cluster. c. Comparison of pairwise gene-gene distances between exact GSPA and fast GSPA embeddings. d. Coexpression networks of top localized genes in each gene cluster. e. *Klf2* KO network.

**Extended Data Figure 12.**
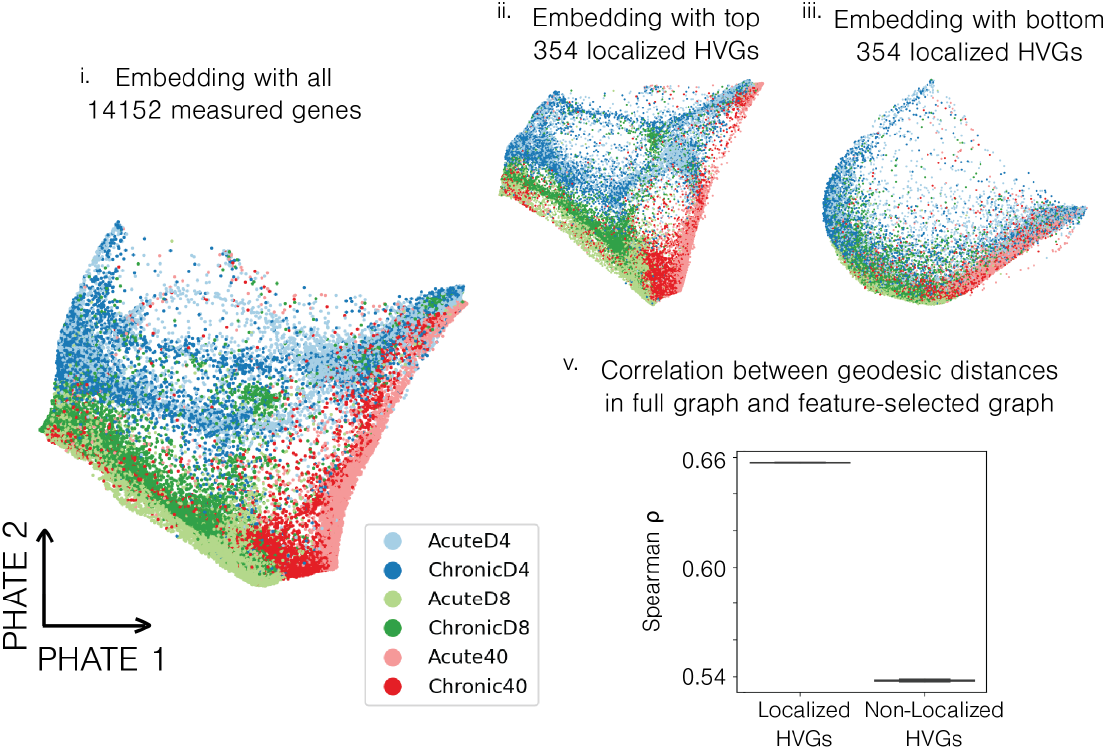
GSPA captures biologically-relevant gene patterns from data. Cell embeddings visualized with PHATE, with (i) all genes (ii) top 25% localized highly variable genes, and (iii) bottom 25% localized highly variable genes. (v). Spearman correlation between random subset of 100,000 pairwise geodesic distances (chosen in two runs) in full cell-cell graph and feature-selected cell-cell graphs.

**Extended Data Figure 13.**
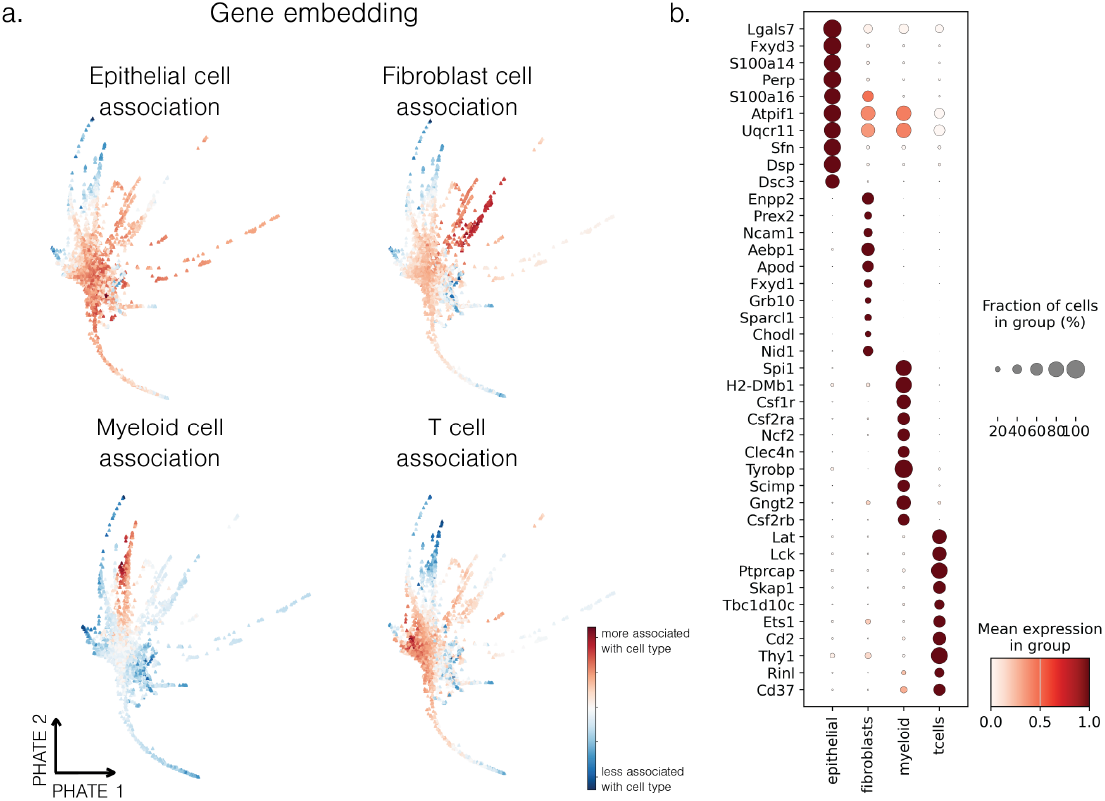
Cell type association scores for four cell types in peripheral tolerance model. a. Gene embedding colored by cell type association ranking. b. Dot plot with top 10 genes associated with each cell type.

**Extended Data Figure 14.**
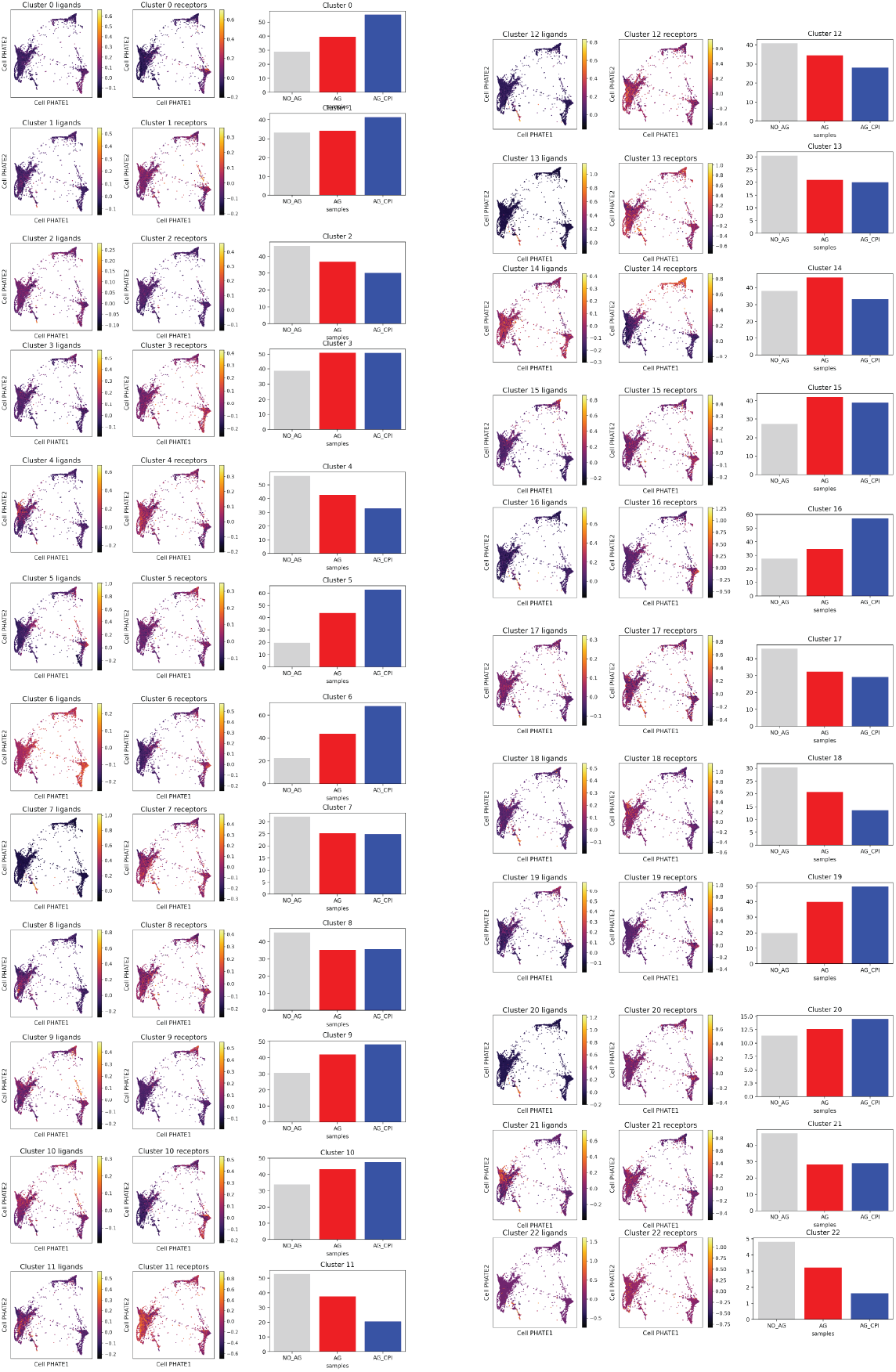
Ligand enrichment score, receptor enrichment score, and enrichment in each condition for all LR pair modules.

**Extended Data Figure 15.**
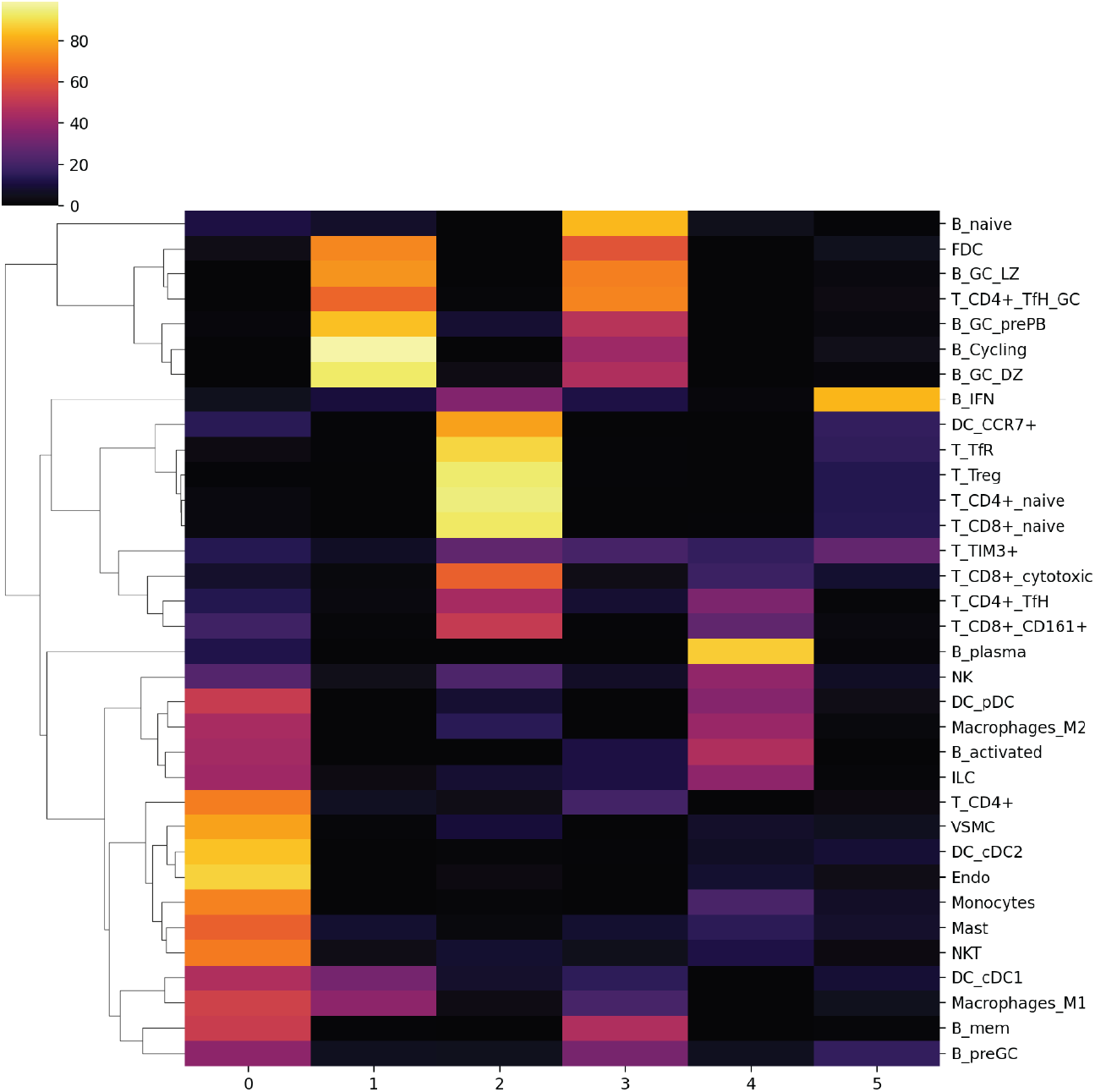
Differentially expressed genes per cell state for each gene module.

